# Defining a highly conserved B cell epitope in the receptor binding motif of SARS-CoV-2 spike glycoprotein

**DOI:** 10.1101/2024.12.06.625234

**Authors:** Sameer Kumar Malladi, Deepika Jaiswal, Baoling Ying, Wafaa B. Alsoussi, Tamarand L. Darling, Bernadeta Dadonaite, Alesandro Civljak, Stephen C. Horvath, Julian Q. Zhou, Wooseob Kim, Jackson S. Turner, Aaron J. Schmitz, Fangjie Han, Suzanne M. Scheaffer, Christopher W. Farnsworth, Raffael Nachbagauer, Biliana Nestorova, Spyros Chalkias, Michael K. Klebert, Darin K. Edwards, Robert Paris, Benjamin S. Strnad, William D. Middleton, Jane A. O’Halloran, Rachel M. Presti, Jesse D. Bloom, Adrianus C. M. Boon, Michael S. Diamond, Goran Bajic, Ali H. Ellebedy

**Author notes:** Contributed equally (Sameer Kumar Malladi, Deepika Jaiswal, Baoling Ying).

## Abstract

SARS-CoV-2 mRNA vaccines induce robust and persistent germinal centre (GC) B cell responses in humans. It remains unclear how the continuous evolution of the virus impacts the breadth of the induced GC B cell response. Using ultrasound-guided fine needle aspiration, we examined draining lymph nodes of nine healthy adults following bivalent booster immunization. We show that 77.8% of the B cell clones in the GC expressed as representative monoclonal antibodies recognized the spike protein, with a third (37.8%) of these targeting the receptor binding domain (RBD). Strikingly, only one RBD-targeting mAb, mAb-52, neutralized all tested SARS-CoV-2 strains, including the recent KP.2 variant. mAb-52 utilizes the IGHV3-66 public clonotype, protects hamsters challenged against the EG.5.1 variant and targets the class I/II RBD epitope, closely mimicking the binding footprint of ACE2. Finally, we show that the remarkable breadth of mAb-52 is due to the somatic hypermutations accumulated within vaccine-induced GC reaction.

**One Sentence Summary:** Booster SARS-CoV-2 mRNA vaccine recruits and broadens GC B cell responses targeting a highly conserved site on receptor binding domain of spike glycoprotein.

## Introduction

The continuous evolution of severe acute respiratory syndrome coronavirus 2 (SARS-CoV-2) has led to the emergence of viral variants of concern and a reduction in the effectiveness of spike (S) protein-derived mRNA vaccines (*1–23*). Multiple studies have demonstrated a reduction in neutralization potency of variants of concern by sera from vaccinees following immunization with a two dose primary series (ancestral WA1/2020) and subsequent boosters (*1, 3–5, 10, 11, 17, 18, 20–22, 24–30*). This result has necessitated the annual update of variant derived vaccines to combat the rise in infections (*31*). Thus, it is imperative to understand whether these variant vaccines primarily induce recall immune responses to the ancestral virus or predominantly *de novo* responses specific to the variants of the vaccine formulation to gauge their effectiveness. Several studies have highlighted the recall responses to ancestral WA1/2020 S protein following variant-derived booster S protein mRNA vaccination (*11, 28, 32–37*). We have previously shown that variant-derived bivalent mRNA-1273.213 (B.1.351/B.1.617.2) or monovalent mRNA-1273.529 (Omicron, BA.1) vaccination in humans predominantly elicited recall memory B cell responses in peripheral blood (*38*), although limited *de novo* memory B cell responses targeting novel epitopes can be detected (*38*).

Vaccination studies predominantly focus on monitoring immune responses in peripheral blood. However, using ultrasound-guided fine needle aspirations (FNAs), it is possible to sample germinal centre (GC) responses in the draining lymph nodes (LNs) (*39–43*) . The GC reaction is responsible for the selection and affinity maturation of antigen-specific B cell clones (*44*), making it critical to understand whether variant-derived vaccines favor the recruitment of memory B cells or naïve clones to induced GCs. Previous studies from our laboratory and others have demonstrated that SARS-CoV-2 mRNA vaccines elicit antibodies targeting three domains: the N-terminal domain (NTD), receptor binding domain (RBD), and S2 domain of the S protein (*40, 45*). The majority of neutralizing antibody responses target the RBD (*46, 47*). RBD-binding antibodies are subclassified into five groups (class I-V) based on their epitope, with class I/II antibodies targeting the receptor binding motif (RBM), neutralizing the virus by blocking its binding to the host receptor, angiotensin-converting enzyme 2 (ACE2) (*46–49*). Antibodies utilizing germline heavy chain genes *IGHV3-53/3-66* and targeting class I/II site have been identified as public clonotypes. Eliciting antibody responses incorporating such clonotypes by vaccination will likely not only result in effective neutralization (*46–48, 50*), but also create a strong selection pressure for population-level escape.

To investigate the impact of SARS-CoV-2 variant booster vaccines on B cell clonal dynamics, GC recruitment and RBD-binding breadth, we enrolled nine healthy participants who had previously received three doses of mRNA-1273 (100 µg)/BNT162b2 (30 µg) into an observational study. Recruited participants received a single (4^th^) dose of 50 µg bivalent mRNA-1273.214 (WA1:BA.1 = 1:1) booster vaccination encoding ancestral and variant-derived S proteins. We used peripheral blood and lymph node FNA samples to assess the degree to which bivalent booster vaccines induced recall or *de novo* B cell responses.

## Results

### B cell responses to mRNA-1273.214 bivalent booster vaccination

Nine participants were recruited to study WU382 in the spring of 2022. After having received three prior mRNA-1273 or BNT162b2 immunizations targeting the WA1/2020 S protein, individuals were boosted with 50 µg of mRNA (1273.214) encoding prefusion stabilized WA1/2020 and BA.1 (97.4% S protein and 93.3% RBD sequence conservation to WA1) S proteins (**Supplementary Data Table S1**). Blood samples were collected at baseline and at weeks 1, 4, 8, and 17 post boosting, and FNAs of draining axillary LNs were collected at the 8-week time point (**Fig. 1A**).

**Figure 1.**
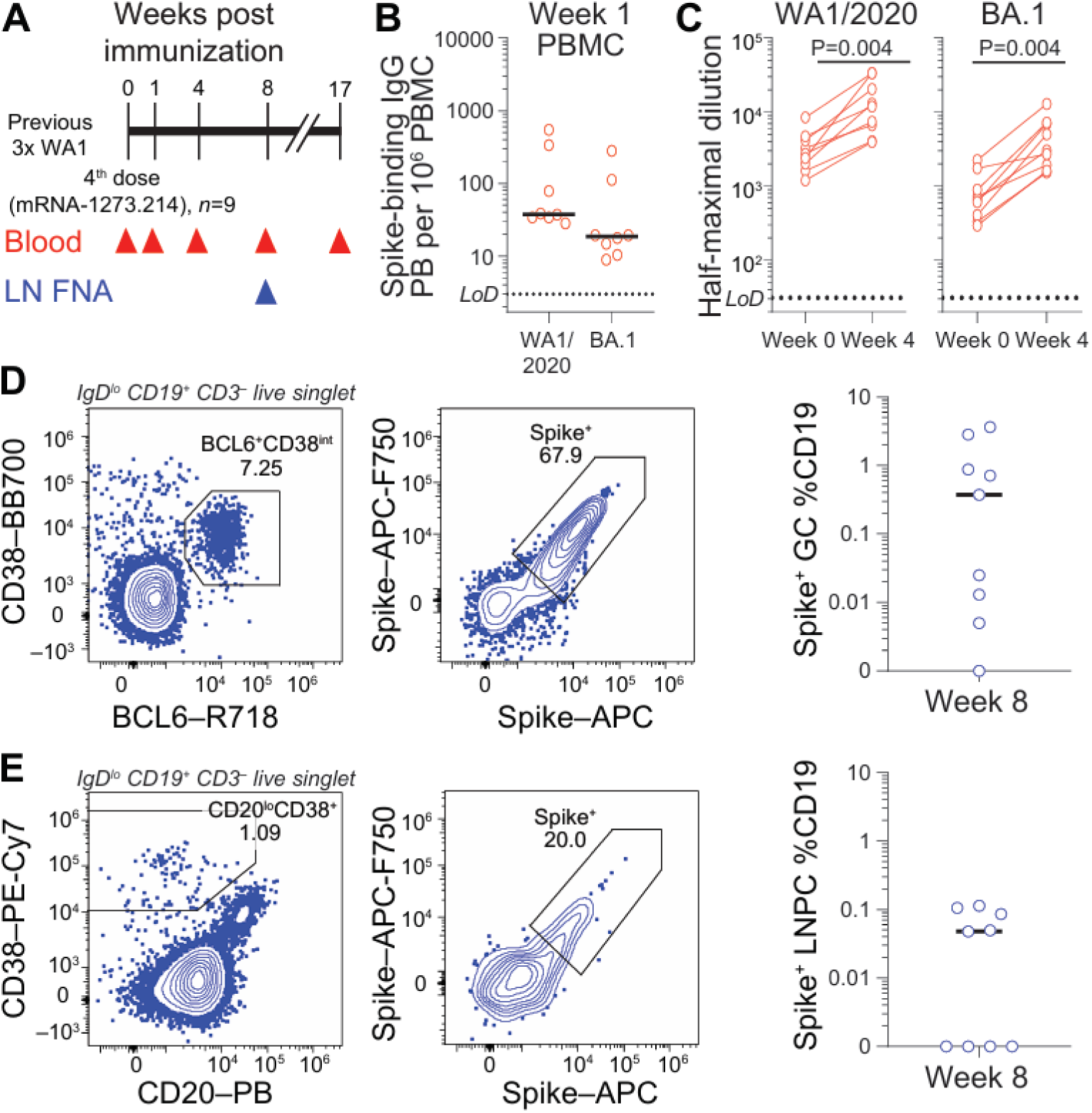
mRNA1273.214 bivalent booster vaccinees’ B cell responses. **(A)** Nine participants immunized previously with three doses of ancestral WA1 vaccine were enrolled and followed after boosting. All participants’ blood was collected at baseline, week 1, 4, 8 and 17 following boosting. Fine needle aspirate of draining axillary lymph nodes were collected from all participants at week 8 post-boosting. (**B)** Frequency of bivalent vaccine WA1 and BA.1 S^+^ antibody responses were probed by ELISpot at week 1 corresponding to peak plasmablast response time-point. (**C)** All participants’ (n = 9) plasma anti-S IgG titres were measured at week 0 and 4 against WA1 and BA.1 Spike protein. Results are from technical duplicates of one experiment. *P* values were determined by two-tailed Wilcoxon matched-pairs signed rank test. (**D)** and **(E)** Representative flow cytometry plots and frequencies of S-binding germinal centre B cells (*BCL6*^+^CD38^int^IgD^lo^CD19^+^CD3^-^) and lymph node plasma cells (CD20^lo^CD38^+^IgD^lo^CD19^+^CD3^-^) of fine needle aspirates from draining axillary lymph nodes at week 8 post boosting. Horizontal lines indicate median in frequency plots.

S^+^ plasmablast (PB) responses were measured in peripheral blood by enzyme-linked immunosorbent spot (ELISpot) assay. We detected robust WA1 and BA.1 S^+^ PB responses one week post boosting in all immunized participants except 382-69, who did not provide sample at day 8 (WA1: 28-547, BA.1: 8-280 S^+^ IgG PBs per million PBMCs) (**Fig. 1B**). The plasma antibody titers increased 2 to 14-fold (Geometric mean titer (GMT): 3.8-fold) against WA1/2020 and 1.5 to 12.3-fold (GMT: 5.3-fold) against BA.1 by week 4 post boosting (**Fig. 1C**). Eight weeks post boosting, draining lateral axillary LNs were sampled by ultrasound-guided FNAs. WA1 and BA.1 S^+^ GC B cells (CD19^+^CD3^-^IgD^lo^Bcl6^+^CD38^int^) were detected in 5 of 9 participants at > 0.1% of CD19^+^ cells (**Fig. 1D**).

### Bivalent boosting recruits extensively cross-neutralizing clones into germinal centres

The FNA samples from the 5 participants with detectable S^+^ GC B cells (382-65/67/69/70/71) were selected for single cell RNA sequencing (scRNA-seq) to track clonal dynamics and determine the antigen specificity of B cell clones (**Fig. 1D-E, Supplementary Fig. S1, S2**). Based on their gene expression profile, scRNA-seq revealed 6 major immune cell clusters typical of a secondary lymphoid organ including B cells, CD4^+^ T cells, CD8^+^ T cells, natural killer cells, monocytes, and plasmacytoid dendritic cells (**Supplementary Fig. S2A, S2B**). Further clustering of B cells (n = 23,128) produced four major subclusters: naïve B cells, germinal centre (GC) B cells, lymph node plasma cells (LNPC) and memory B cells (MBC) (**Fig. 2A, Supplementary Fig. S2C, S2D**). Using paired heavy and light chain B cell receptor (BCR) sequencing data, we computationally recovered 598 clonally distinct GC B cell and LNPC clones for monoclonal antibody (mAb) generation. We characterized the S protein binding of these mAbs by enzyme-linked immunosorbent assay (ELISA) and mapped the S^+^ mAbs to B cell clones consisting of 2086 single cells (n = 2086) across multiple B cell subclusters (**Fig. 2A, right panel**). A major fraction (n = 465, 77.8%) of the mAbs were WA1 S^+^, while the remaining 22.2% (n = 133) clones were non-S-binders (**Fig. 2B**). The entirety of the S^+^-binders bound WA1, suggesting the GC response was due to recall of MBCs previously exposed to the ancestral S antigen (**Fig. 2A-B, Supplementary Fig. S2E**). Further, we longitudinally tracked S-binding GC B cell clones in peripheral blood at baseline (MBC), week 1 (PB), and week 17 (MBC). Clonal tracking revealed consistently high percentages of clonal overlap of GC B cells to PB at week 1, and less consistently so of GC B cells to MBC at baseline, further confirming that the GC B cell response was recall-derived (**Supplementary Fig. S3**). We further determined if the S-binders specifically targeted the major neutralizing domain, the RBD. Of the S-specific mAbs, 37.8% (n = 176) bound ancestral WA1- derived RBD (RBD^+^), while the remaining were non-RBD (RBD^-^) binders (**Fig. 2C**).

**Figure 2.**
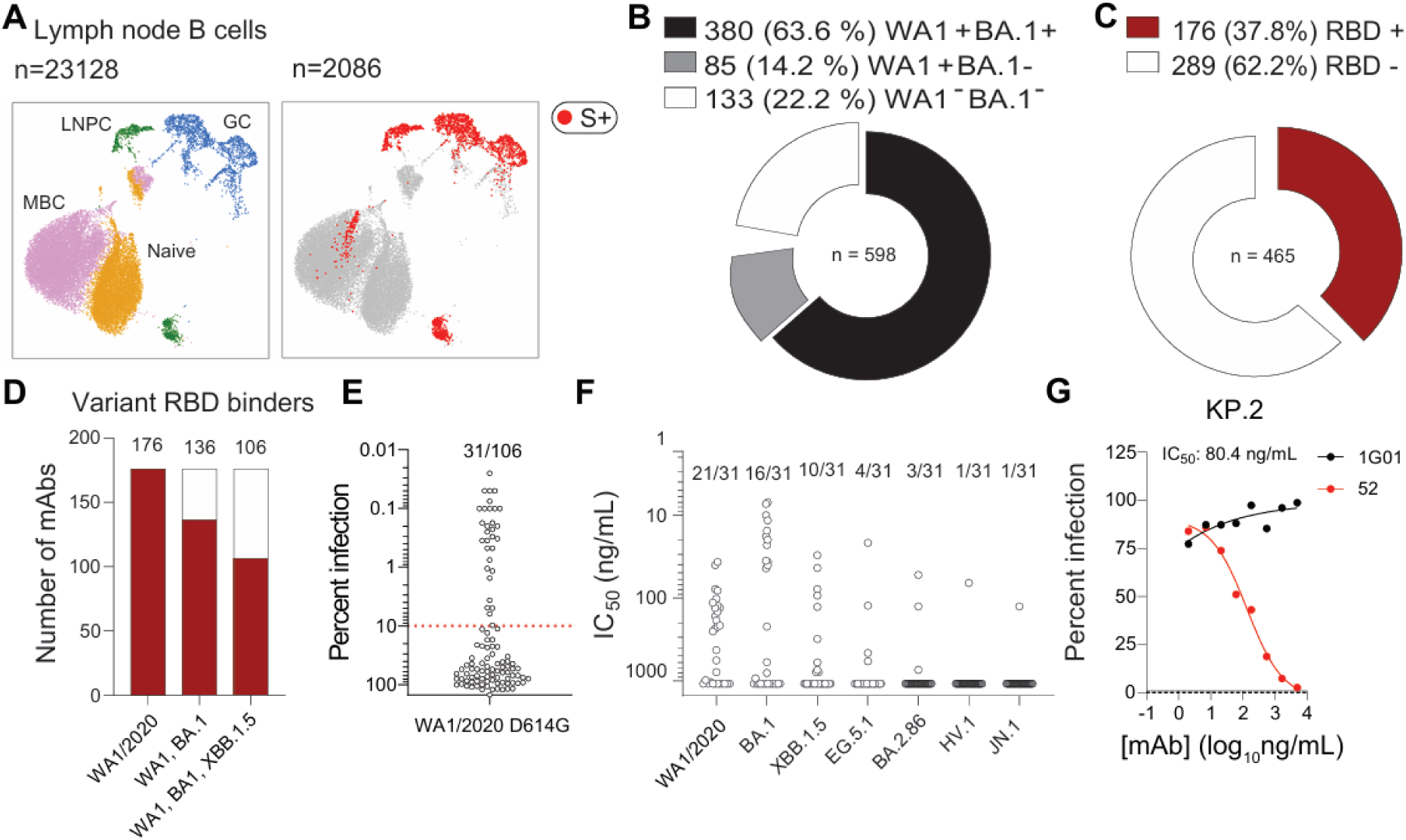
Lymph node fine needle aspirate B cell analysis of vaccinees. **(A)** Uniform manifold approximation and projection (UMAP) representing B cell transcriptional clusters from single cell RNA sequencing (scRNA-seq) of lymph node FNAs (left) with spike-specific clones overlaid (right). Each dot represents a cell, coloured by phenotype as defined by transcriptomic profile (left) and S-specificity (right). Total number of cells (left) and number of S^+^ cells (right) are on the top left corner. GC, GC B cell; LNPC, Lymph node plasma cell; MBC, Memory B cell. (**B)** Antigen binding landscape of GC B cell and LNPC mAbs (n = 598) derived from distinct B cell clonal lineages from lymph node FNA at week 8 of five participants. (**C)** Binding of mAbs from S- specific GC B cells and LNPCs at day 57 post boosting to constituent domains of WA1/2020 S protein measured by ELISA. (**D)** WA1/2020 RBD-binding mAbs binding variants of concern RBD. The number of mAbs binding RBD of WA1/2020; WA1/2020 and BA.1; and WA1/2020, BA.1, and XBB.1.5 are depicted. (**E)** Neutralizing activity of mAbs cross reacting with WA1/2020, BA.1, and XBB.1.5 as determined with VSV-S (WA1/2020 D614G) chimeric virus assays. Each symbol represents one mAb. Percentages indicate the proportion of mAbs above 90% infection reduction threshold. **(F)** Neutralizing activity of mAbs in infection against WA1/2020 D614G, BA.1, XBB.1.5, EG.5.1, BA.2.86, HV.1, JN.1 **(F)** and KP.2 **(G)** authentic viruses. Each symbol in **(F)** represents one mAb. Authentic virus neutralization IC50 quantitated in ng/ml and mAbs are considered neutralizing given IC50 < 1000 ng/ml. mAb 52 potently neutralized WA1 D614G, BA.1, XBB.1.5, EG.5.1, BA.2.86, HV.1, JN.1 **(F)** and KP.2 **(G)** viral variants. Results are from technical duplicates of one experiment.

We next characterized the cross reactivity of the clones that bound WA1 RBD (n = 176). A majority (60.2%) of the WA1 RBD-binding mAbs cross reacted with RBDs from both BA.1 and XBB.1.5, a subsequent variant of concern (n = 106) (**Fig. 2D**). A minority (29.2%) of these cross reactive mAbs neutralized chimeric vesicular stomatitis viruses (VSV) expressing the WA1/2020 D614G S protein (n = 31, > 90% inhibition) in a single endpoint neutralization (10 µg/ml) (**Fig. 2E**). We further assessed the neutralization capacity of these mAbs using an authentic virus neutralization assay against WA1, BA.1, XBB.1.5, EG.5.1, BA.2.86, HV.1, and JN.1 variants (**Fig. 2F**). We observed a gradual decrease in the number of mAbs that retained inhibitory activity as the antigenic distance of the virus increased from the ancestral strain: WA1 (n = 21); BA.1 (n = 16); XBB.1.5 (n = 10); EG.5.1 (n = 4); BA.2.86 (n = 3); HV.1 (n = 1); and JN.1 (n = 1) (**Fig. 2F**).

A minority (n = 10) of the mAbs that neutralized VSV-WA1/2020 D614G S at high concentration lost the ability to inhibit infection of authentic WA1/2020 virus, possibly due to differences in the virus neutralization assays, concentration of tested mAbs (10 µg/ml vs 5 µg/ml), or relative expression of S protein on the virion. Notably, we observed a single mAb, mAb-52, which potently neutralized all tested viral strains: ancestral WA1 (IC50: 51.9 ng/mL) and variants BA.1 (IC50: 5.5 ng/mL), XBB.1.5 (IC50: 12.4 ng/mL), EG.5.1 (IC50: 16.2 ng/mL), BA.2.86 (IC50: 42.1 ng/mL), HV.1 (IC50: 53.3 ng/mL), JN.1 (IC50: 60.6 ng/mL), and KP.2 (IC50: 80.4 ng/mL) (**Fig. 2F, 2G, Supplementary Fig. S4**). These results highlight the recruitment of some broadly cross-reactive germinal centre B cell clones following bivalent booster vaccination.

### GC B cell-derived mAb-52 protects hamsters from EG.5.1 challenge

Given the breadth of mAb-52 and possible treatment implications, we next examined its protective efficacy against the EG.5.1 SARS-CoV-2 variant in a hamster challenge model. Six hamsters were treated with mAb-52 or isotype control antibody 1G05 (dose: 10 mg/kg) one day prior to intranasal challenge with the EG.5.1 (10^4^ PFU) variant of SARS-CoV-2 (**Fig. 3A**). Following challenge, the hamsters were monitored for three days prior to measurement of infectious virus and viral RNA in the nasal wash, nasal turbinate, and left lung homogenates (**Fig. 3A**). mAb-52-treated hamsters had lower levels of infectious virus in the upper and lower respiratory tract compared to isotype control-treated hamsters (nasal wash: 23-fold, nasal turbinate: 103-fold, lungs:11,195-fold) (**Fig. 3B-D, left panels**). Similarly, we observed lower levels of viral RNA in mAb-52-treated hamsters than the isotype control-treated animals (nasal wash: 8.6-fold, nasal turbinate: 4.7-fold, lungs: 1,804-fold) (**Fig. 3B-D, right**).

**Figure 3.**
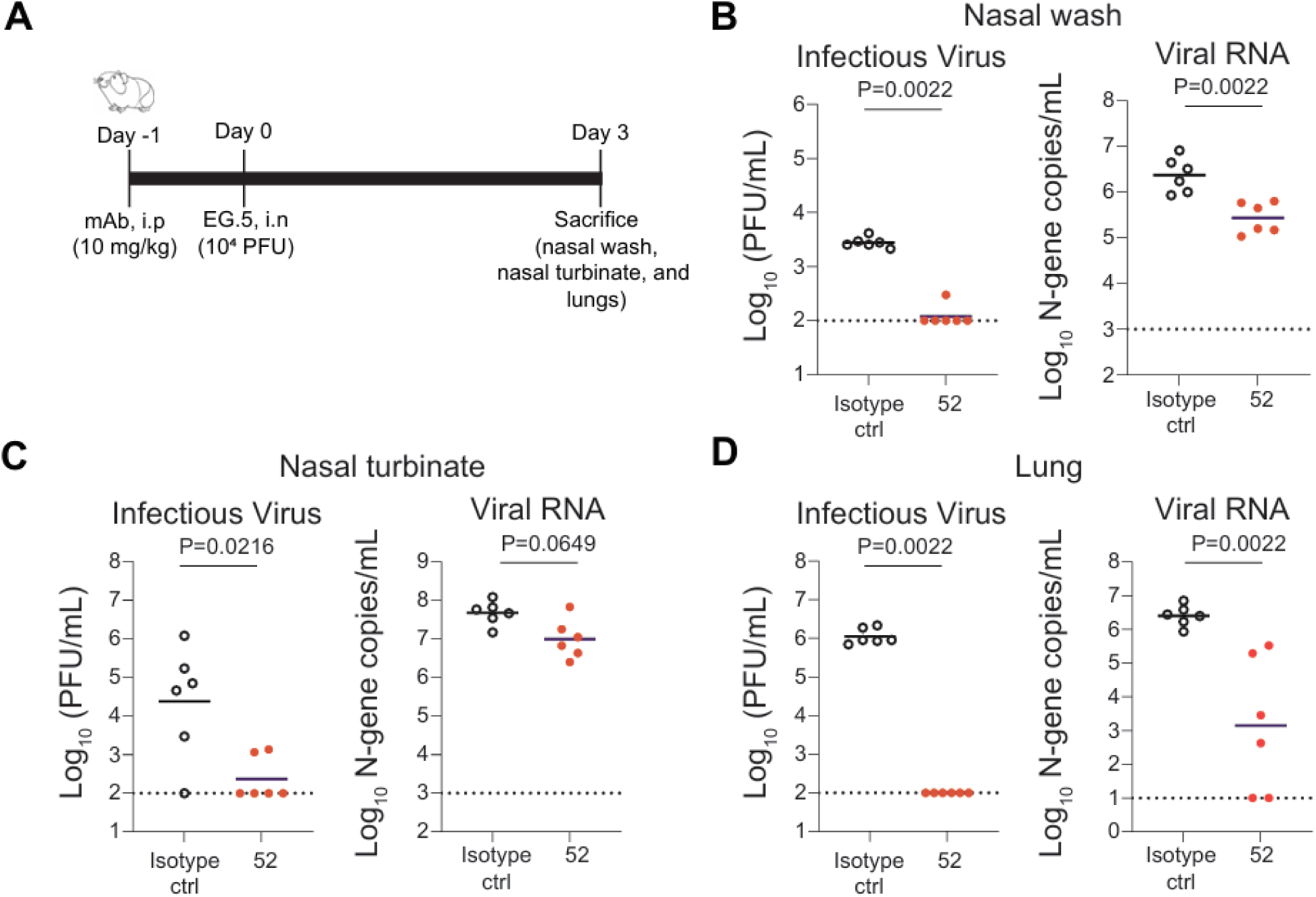
mAb-52 protects hamsters from EG.5.1 challenge. (**A**) EG.5.1 challenge of five- week-old male Syrian golden hamsters. One day prior to challenge (d-1), the hamsters received an intraperitoneal injection with mAb-52 or 1G05 isotype control at 10 mg/kg. The following day (d0), the hamsters were intranasally challenged with 10^4^ PFU of SARS-CoV-2 EG.5.1. Following challenge, the hamsters were monitored daily and their (**B**) nasal wash, (**C**) nasal turbinate, and (**D**) left lungs were harvested on day 3 for measurement of infectious virus by plaque assay and viral RNA by RT-qPCR. The data is from one experiment with 6 hamsters per group/experiment. *P* values were determined by two-tailed Mann-Whitney test.

### mAb-52 belongs to public clonotype IGHV3-66*02 and targets the class I/II RBD epitope

We observed several public B cell clonotypes expressed in the FNA of vaccinated individuals comprising IGHV1-69/3-23/3-30/3-33/4-39/5-51 (**Supplementary Fig. S2F**). mAb-52 is encoded by a frequently utilized public clonotype, IGHV3-66*02 (*48, 51*). Antibodies utilizing germline heavy chain gene IGHV3-66 and binding the RBD often target class I/II epitopes on the RBD (*48, 51*). Consistent with prior observations (*47, 48, 52*), mAb-52 bound with nanomolar affinity to all RBD variants tested: WA1 (KD = 0.6 nM), BA.1 (KD = 1.1 nM), EG.5.1 (KD = 7 nM), HV.1 (KD = 9.1 nM), JN.1 (KD = 23.8 nM), and KP.2 (KD = 139 nM) (**Supplementary Fig. S5A**). We note that the binding affinity is decreasing against newer variants, even though it remains high. mAb- 52 competed with ACE2 for binding the RBD, as well as with a previously characterized class I/A-binding monoclonal antibody (2B04) (*53*), suggesting mAb-52 might engage an epitope similar to or near the ACE2 receptor binding site (**Supplementary Fig. S5B**).

We employed pseudovirus deep mutational scanning (DMS) using libraries of the XBB.1.5 spike with saturating RBD mutations (*54*) to identify the key sites where mutations escape neutralization by mAb-52. Escape from mAb-52 is caused primarily by mutations at nine spatially clustered sites (420, 421, 455, 456, 473, 475, 487, 488, and 491) (**Fig. 4A**). At many of these sites, some amino-acid mutations cause more escape than others (**Fig. 4B**). For instance, at site L455, mutations to charged or large amino acids (eg, L455D, L455E and L455W) caused strong escape; but mutations to some other amino acids had only a modest effect, likely explaining why mAb-52 still neutralized JN.1 (which contains L455S). We validated key escape mutants by two independent *in vitro* assay platforms: ELISA testing mAb-52, and biolayer interferometry (BLI) testing the fragment antigen binding (Fab) of mAb-52 (**Fig. 4C, Supplementary Fig. S6A-C**). For these validation experiments, escape mutants were incorporated in the background of ancestral RBD as majority of the escape mutant positions were conserved across XBB.1.5 and WA1/2020. ELISA and BLI assays corroborated the loss of binding, with escape residues losing binding (>10-percent decrease in ELISA AUC or >10-fold increase in BLI KD) to mAb-52 and Fab-52, respectively (**Fig. 4C-D, Supplementary Fig. S6A-C**). Key escape sites determined by DMS for which we validated the effects of mutations in these *in vitro* assays included residues 420, 421, 455, 456, 473, 475, 487 and 491 (**Fig. 4C-D**).

**Figure 4.**
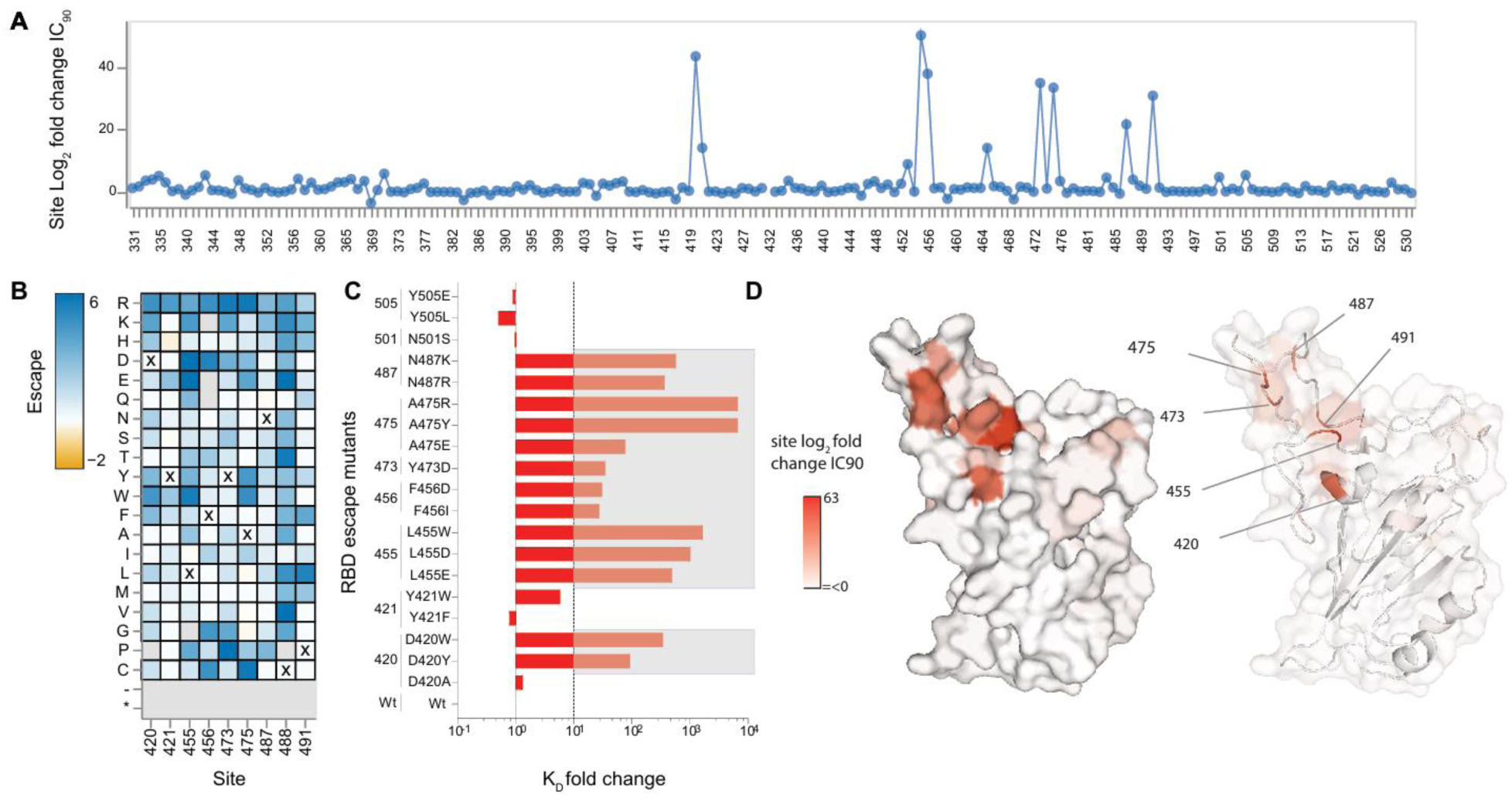
XBB.1.5 RBD deep mutational scanning escape of mAb-52. **(A)** Total escape at each site in the XBB.1.5 RBD as measured by pseudovirus deep mutational scanning library. (**B)** Escape caused by individual mutations at key sites of escape. XBB.1.5 wild type amino acids are depicted with X and amino acids in grey are absent in the library or highly deleterious for spike function. The complete data for line plot, heat map and analysis code can be accessed at dms-vep.org/SARS-CoV-2_XBB.1.5_RBD_DMS_mAB-52/htmls/mAb_52_mut_icXX.html **(C)** KD fold change determined by BLI binding of RBD escape mutants binding Fab-52. RBD mutants with KD fold change >10 are considered the footprint of mAb-52. **(D)** Mutational escape at key residues highlighted in heat map mapped onto surface representation of RBD. The higher intensity depicts higher escape. The epitope can be categorized as class I/II based on previous RBD antibody nomenclature. Results are the average of two independent libraries.

### Cryo-EM structure of mAb-52 complexed with XBB.1.5 spike

To gain greater molecular insight into the binding epitope targeted by mAb-52, we determined the structure of XBB.1.5 S protein in complex with Fab-52 by cryo-electron microscopy (cryo-EM), yielding a global resolution map at 2.6Å (**Fig. 5A-B, Supplementary Fig. S7A-E**). The XBB.1.5 S used for cryo-EM contains FLip mutations in the RBD (L455F and F456L). Fab-52 bound to one RBD in up-conformation on S protein trimer (**Fig. 5A**). We improved the antibody:antigen interface resolution by local refinement to achieve a map at 3.1 Å of nominal resolution (**Fig. 5B, Supplementary Fig. S7E**). The cryo-EM structure confirmed that mAb-52 targets the class I/II RBD epitope, with the key binding residues in the receptor binding groove composed of F455, K460, Y473, V475, N477, N487, Q493 and the receptor binding motif (RBM) lateral residues D420, Y421 (**Fig. 5C-D, Supplementary Fig. S8**). The Fab heavy chain bound the residues in the RBM and the lateral surface, which were otherwise occluded on the RBD in the down conformation. mAb-52 targets several secondary structural regions of the RBD, including alpha helices α4 and α5, and beta sheets β5 and β6 in the RBM, and alpha helix α3 in the periphery of the RBM. The light chain provides supporting interactions with residues R403, G502, V503, and H505 (**Fig. 5E-F**). The approach angle of mAb-52 binding XBB.1.5 RBD is consistent with previously reported RBD-binding mAbs utilizing the IGHV3-66*02 public clonotype, including CS23, PDI37, C98C7, and others (*55*). There are several antibodies characterized with similar HC:LC pairing as mAb-52, but none with determined structures (*55*). mAb-52 complexed with S protein cumulatively buried a surface area of 1190 Å^2^ with a higher contribution from the heavy chain (804 Å^2^) and lower contribution from the light chain (386 Å^2^) (**Supplementary Fig. S9A- B**). Fab-52 engages RBD via polar and hydrophobic interactions involving CDRH1-H3, CDRL1 and CDRL3. mAb-52 residue Y33 of the CDRH1 formed H-bonding interactions with F455 backbone carbonyl oxygen (**Fig. 5E-F**). G54 and S56 within the CDRH2 formed extensive H-bonding interactions with terminal carbonyl oxygen atoms of Y421 and D420, respectively. Additionally, R97 and E101 of the CDRH3 formed H-bonding with RBD residues N487 and Q493 respectively. Notably, of the 15 hydrogen bonds between Fab-52 and the RBD, six involve the RBD backbone rather than the amino acid side chains (specifically, residues F455, V475, N477, G502 and V503), which allow mAb-52 to maintain its effectiveness despite mutations in the virus spike glycoprotein.

**Figure 5.**
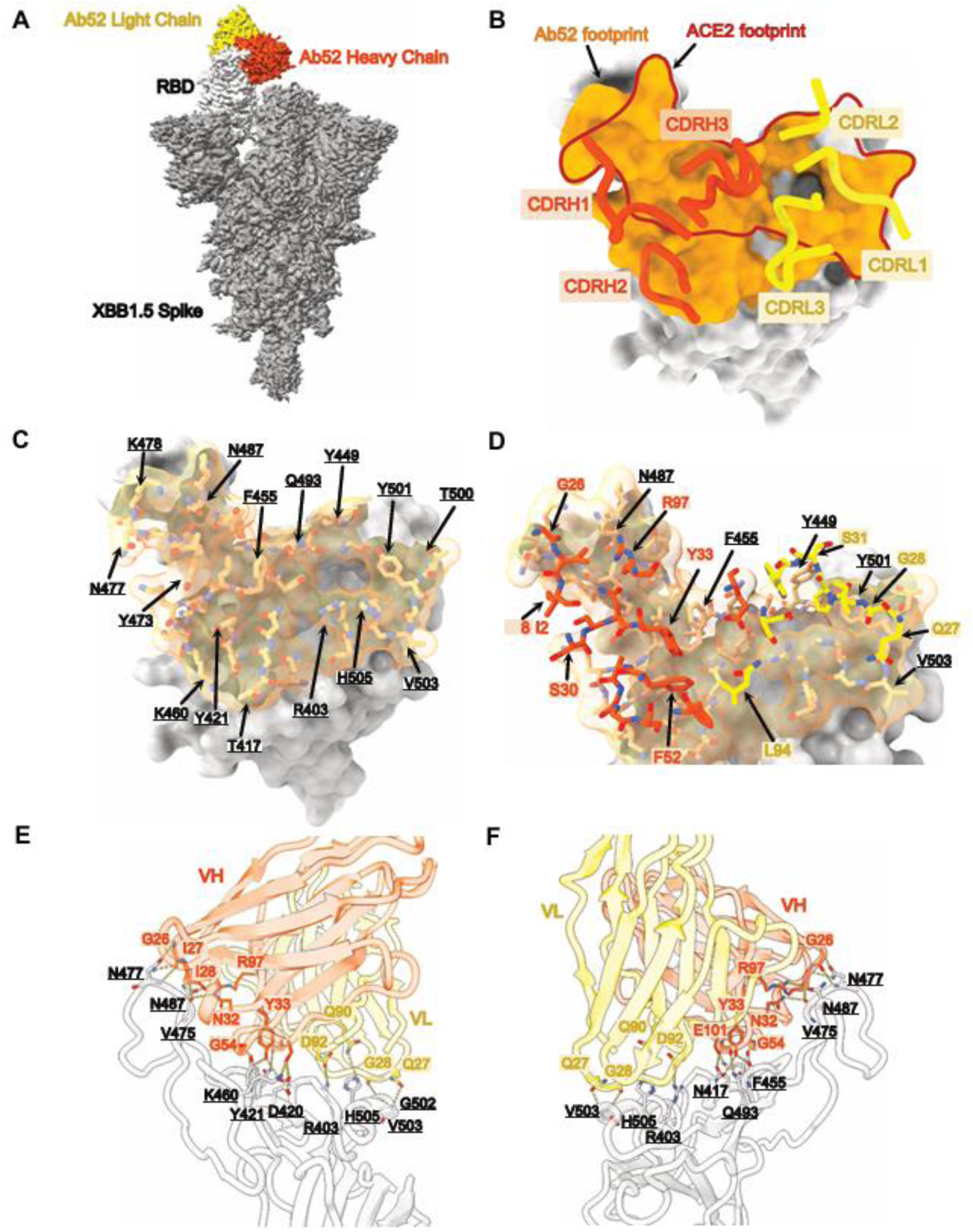
Cryo-EM structure of Fab-52 in complex with XBB.1.5 spike. **(A)** 2.58 Å cryo-EM density for Fab-52-XBB.1.5 Spike trimer complex; Fab-52 VH (orange red) and VL (yellow) bound to the RBD (light gray). (**B)** Fab-52 footprint (orange), as defined by buried surface area, depicted on a surface representation of the RBD (light gray) with CDR loops of the Fab-52 VH (orange red) and VL (yellow), ACE2 footprint is outlined as red line track on RBD. (**C)** RBD epitope residues denoted by arrows in FAb-52:RBD interface. (**D)** Fab-52 paratope residues denoted by arrows in orange red and yellow. (**E) and (F)** are two 180° views along the y-axis that show details of the Fab-52:RBD molecular interface with numerous polar interactions.

Previous work indicates somatic hypermutations F28I and Y57F in the IGHV3-66/IGHV3-53 public clonotypes led to affinity maturation and Omicron RBD binding (*52, 56*). However, we did not observe side chains of these amino acids interacting with RBD, and reversion of these amino acids to germline (I28F and F57Y) did not lead to a reduction or abrogation of binding to BA.1 Omicron RBD (**Supplementary Fig. S10A**). We determined the binding affinity of the mature and germline Fab-52 to variant RBD proteins. The mature Fab-52 bound all the variant RBD proteins in nanomolar affinity from highest to lowest order as WA1/ BA.1 (0.6 nM) > EG.5.1 (10 nM) > HV.1 (12.2 nM) > JN.1 (28 nM) > KP.2 (139 nM) (**Supplementary Fig. S10B**). Germline Fab-52 bound WA1 (1.9 nM) and BA.1 (29.3 nM) RBD with nanomolar affinity, but lost binding to later Omicron variants EG.5.1, HV.1, JN.1 and, KP.2 (**Supplementary Fig. S10B**). These results highlight the recruitment of some broadly cross-reactive, protective, and public B cell clones targeting class I/II RBD epitope to the germinal centre following bivalent booster vaccination.

## Discussion

We evaluated the GC B cell response to a bivalent (mRNA1273.214) SARS-CoV-2 vaccine booster in humans. All the participants responded to bivalent mRNA-1273.214 boosting based on frequencies of antibody-secreting S^+^ PBs in blood and increased serum binding titres to S protein (WA1/2020, BA.1). We observed a robust S^+^ GC response in the majority of vaccinees upon boosting with the bivalent vaccine. All the S^+^ GC B cell clones bound ancestral S-protein, highlighting potential recall responses recruited to the GC with no or very limited variant-specific naïve responses, possibly due to inclusion of the ancestral S protein (WA1/2020)-encoding mRNA to the bivalent booster. A major fraction (60.2 %) of the RBD-targeting clones were extensively cross-reactive (WA1^+^BA.1^+^XBB.1.5^+^) with a single clone encoding mAb-52, which potently cross-neutralized newer Omicron variants through KP.2. mAb-52 targeted the class I/II RBD epitope, utilized a public clonotype IGHV3-66, and protected hamsters challenged with a distant EG.5.1 variant. These results highlight that bivalent booster vaccination can recruit some broadly neutralizing, public clones, indicating wider applicability and success at eliciting such responses in vaccinees. We note that the observed mAb-52 response was rare due to limited number of clones analysed from a single time point FNA. Taken together, booster vaccination with variant-derived S-encoding mRNA could broaden the elicited responses by minimizing recruitment of naïve ancestral (WA1) S-binding clones. Furthermore, it is not surprising to observe a dominant antigenically imprinted serological responses as reported by several studies (*11, 28, 30, 32, 33, 35–37, 57, 58*), owing to the addition of ancestral S-encoding mRNA to the vaccine. Nonetheless, this type of imprinting effect was seen previously with serum antibodies that gained cross-reactive neutralizing activity against distantly related Sarbecoviruses after boosting monovalently with BA.5 or XBB.1.5 mRNA vaccines (*35*). While a recent study indicated bivalent vaccination led to elicitation of heightened IgG4 subclass sera responses, we did not observe such IgG4 skewing of germinal centre B cell responses at week 8 in all our participants (*58*). Another study recently reported that the JN.1 variant evades majority of antibodies utilizing IGHV3-53/66, whereas our study shows that mAb-52, also derived from IGHV3-66, broadly neutralizes evolved variants of concern, JN.1 and KP.2 (*46–48, 59*). These results suggest an affinity matured public clonotype is amenable to recruitment and further affinity maturation to neutralize Omicron-lineage descendants that evolve in the future. Eliciting such broad cross-neutralizing antibody responses like mAb-52 is the primary goal of a variant-derived vaccination that we could successfully capture in the GC B cell compartment of the lymph nodes following bivalent SARS-CoV-2 booster vaccination.

## Limitations of the study

The number of lymph nodes (n = 1) sampled per participant and lack of longitudinal sampling limits the evaluation of rare and sporadic occurrences of B cell clonal representatives that exclusively bind variant S protein epitopes. Successive FNA sampling of humans receiving divergent variant-derived S protein vaccines would help test the elicitation of variant-derived GC responses. A consideration for future studies is to test the relative expression, immunogenicity, and stability of component S proteins being expressed and how this modulates antibody responses given its wider applicability in case of upcoming combined COVID-Flu and other respiratory virus vaccines (*60–62*). Further, we only tested the neutralizing capacity of RBD-binding mAbs and did not test other potential effector functions such as antibody dependent cellular cytotoxicity (ADCC) or antibody dependent cellular phagocytosis (ADCP), which also can confer protection in the absence of neutralization (*63*).

## Materials and Methods

### Sample collection, preparation, and storage

All studies were approved by the Institutional Review Board of Washington University in St. Louis. Written consent was obtained from all participants. Nine healthy volunteers were enrolled, of whom all provided axillary LN (**Supplementary Data Table S1**). Blood samples were collected in ethylenediaminetetraacetic acid (EDTA) evacuated tubes (BD), and peripheral blood mononuclear cells (PBMC) were enriched by density gradient centrifugation over Lymphopure (BioLegend). The residual red blood cells were lysed with ammonium chloride lysis buffer, washed with PBS supplemented with 2% FBS and 2 mM EDTA (P2), and PBMC were immediately used or cryopreserved in 10% dimethylsulfoxide (DMSO) in FBS. Ultrasound-guided FNA of axillary LNs was performed by a radiologist. LN dimensions and cortical thickness were measured, and the presence and degree of cortical vascularity and location of the LN relative to the axillary vein were determined prior to each FNA. For each FNA sample, six passes were made under continuous real-time ultrasound guidance using 22- or 25-gauge needles, each of which was flushed with 3 mL of RPMI 1640 supplemented with 10% FBS and 100 U/mL penicillin/streptomycin, followed by three 1-mL rinses. Red blood cells were lysed with ammonium chloride buffer (Lonza), washed with P2, and immediately used or cryopreserved in 10% DMSO in FBS. Participants reported no adverse effects from phlebotomies or serial FNAs.

### Cell lines

#### Expi293F cells were cultured in Expi293 Expression Medium (Gibco)

Vero cells expressing human ACE2 and TMPRSS2 (Vero-hACE2-hTMPRSS2) (*64*) were cultured at 37°C in Dulbecco’s Modified Eagle medium (DMEM) supplemented with 10% fetal bovine serum (FBS), 10 mM HEPES (pH 7.3), 100 U/mL of penicillin, 100 µg/ml of streptomycin, and 10 µg/ml of puromycin. Vero cells expressing TMPRSS2 (Vero-hTMPRSS2) were cultured at 37°C in Dulbecco’s Modified Eagle medium (DMEM) supplemented with 10% fetal bovine serum (FBS), 10 mM HEPES (pH 7.3), 100 U/mL of penicillin, 100 µg/ml of streptomycin, and 5 µg/ml of blasticidin.

#### Antigens

Recombinant soluble spike protein (S) from WA1/2020 (2P), B.1.351 (2P), B.1.617.2 (2P), BA.1 (6P) strains of SARS-CoV-2 and their Avi-tagged counterparts were expressed as previously described(*41, 43*). Briefly, mammalian cell codon-optimized nucleotide sequences coding for the soluble ectodomain of S (GenBank: MN908947.3, amino acids 1-1213) including a C-terminal thrombin cleavage site, T4 foldon trimerization domain, and hexahistidine tag (2P version)/octa histag (6P version) were cloned into mammalian expression vector pCAGGS. The S sequences were modified to remove the polybasic cleavage site (RRAR to A in WA1 and RRAR to GSAS in BA.1) and 2P (K986P and V987P) (*65*), 6P (F817P, A892P, A899P, A942P, K986P and V987P) (*66*). For expression of Avi-tagged variants, the CDS of pCAGGS vector containing the sequence for the relevant soluble S was modified to encode 3’ Avitag insert after the HIS tag (5’-HIS tag- GGCTCCGGGCTGAACGACATCTTCGAAGCCCAGAAGATTGAGTGGCATGAG-Stop-3’; HIS tag-GSGLNDIFEAQKIEWHE-Stop). Recombinant proteins were produced in Expi293F cells (ThermoFisher) by transfection with purified DNA using the ExpiFectamine 293 Transfection Kit (ThermoFisher). Supernatants from transfected cells were harvested 3 days post- transfection, and recombinant proteins were purified using Ni-NTA agarose (ThermoFisher), then buffer exchanged into phosphate buffered saline (PBS) and concentrated using Amicon Ultracel centrifugal filters (EMD Millipore). To biotinylate Avi-tagged S variants, the S-Avitag substrates were diluted to 40 μM and incubated for 1 h at 30℃ with 15 µg/ml BirA enzyme (Avidity) in 0.05 M bicine buffer at pH 8.3 supplemented with 10 mM ATP, 10 mM MgOAc, and 50 µM biotin. The protein was then concentrated/buffer exchanged with PBS using a 100 kDa Amicon Ultra centrifugal filter (MilliporeSigma).

To generate antigen probes for flow cytometry staining and sorting, trimeric BirA-biotinylated recombinant S from WA1/2020 or BA.1 (mRNA-1273) were incubated with a 1.04-fold molar excess of BV421-, BV650-, or PE-conjugated streptavidin (BioLegend) on ice, with three equal additions of S spaced every 15 min. Fifteen min after the third S addition, D-biotin was added in 6-fold molar excess to streptavidin to block any unoccupied biotin binding sites. SA-PE-Cy5 was blocked with a 6-fold molar excess of D-biotin and used as a background staining control. Bovine serum albumin (BSA) was biotinylated using the EZ-Link Micro NHS-PEG4-Biotinylation Kit (Thermo Fisher); excess unreacted biotin was removed using 7-kDa Zeba desalting columns (Pierce).

#### ELISpot assay

Wells of a microtiter plate were coated with recombinant S from the WA1/2020, B.1.351, B.1.617.2, BA.1, BSA or pooled anti-κ and anti-λ light chain antibodies (Cellular Technology Limited). Direct *ex-vivo* ELISpot assays were performed to determine the number of total, recombinant S-binding IgG- and IgA-secreting cells present in PBMC and enriched BMPC samples using IgG/IgA double-color ELISpot Kits (Cellular Technology Limited) according to the manufacturer’s instructions. Plates were analyzed using an ELISpot counter (Cellular Technology Limited).

#### ELISA

Assays were performed in 96-well MaxiSorp plates (Thermo Fisher) coated with 100 µL of recombinant SARS-CoV-2 S from WA1/2020 (2P), BA.1 (6P) and RBDs from WA1, BA.1, XBB.1.5 strains of SARS-CoV-2 bovine serum albumin diluted to 1µg/ml in PBS, and plates were incubated at 4°C overnight. Plates then were blocked with 10% FBS and 0.05% Tween 20 in PBS. Plasma or purified monoclonal antibodies were diluted serially starting at 1:30 or at fixed concentration of 10 µg/ml respectively in blocking buffer and added to the plates. Plates were incubated for 90 min at room temperature and then washed 3 times with 0.05% Tween 20 in PBS. Goat anti-human IgG-HRP secondary antibody (goat polyclonal, Jackson ImmunoResearch, 109- 035-088, 1:2,500) was diluted in blocking buffer before adding to plates and incubating for 60 min at room temperature. Plates were washed 3 times with 0.05% Tween 20 in PBS and 3 times with PBS before the addition of o-phenylenediamine dihydrochloride peroxidase substrate (Sigma-Aldrich). Reactions were stopped by the addition of 1 M hydrochloric acid. Optical density were measured at 490 nm.

#### Flow cytometry and cell sorting

Staining for flow cytometry analysis was performed using cryo-preserved FNA samples. For analysis, FNA samples were incubated for 30 min on ice with purified CD16 (3G8, BioLegend, 1:100), CD32 (FUN-2, BioLegend, 1:100), CD64 (10.1, BioLegend, 1:100) and PD-1-BB515 (EH12.1, BD Horizon, 1:100) in P2, washed twice, then stained for 30 min on ice with WA1/2020 probes pre-conjugated to SA-APC and SA-APC-Fire 750, BA.1 probes pre-conjugated to SA- BV421 and SA-BV650, biotin-saturated SA-PE-Cy5, IgG-BV480 (goat polyclonal, Jackson ImmunoResearch, 1:100), IgA-FITC (M24A, Millipore, 1:500), CD8-A532 (RPA-T8, Thermo, 1:100), CD38-BB700 (HIT2, BD Horizon, 1:500), CD20-Pacific Blue (2H7, 1:400), CD4-Spark Violet 538 (SK3, 1:400), IgM-BV605 (MHM-88, 1:100), CD19-BV750 (HIB19, 1:100), IgD-BV785 (IA6-2, 1:200), CXCR5-PE-Dazzle 594 (J252D4, 1:50), CD14-PerCP (HCD14, 1:50), CD71-PE-Cy7 (CY1G4, 1:400), CD27-PE-Fire 810 (O323, 1:200), CD3-APC-Fire 810 (SK7, 1:50), and Zombie NIR (all BioLegend) diluted in Brilliant Staining buffer (BD Horizon). Cells were washed twice with P2, fixed for 1 h at 25°C using the True Nuclear fixation kit (BioLegend), washed twice with True Nuclear Permeabilization/Wash buffer, stained with Ki-67-BV711 (Ki- 67, BioLegend, 1:200), Blimp1-PE (646702, R&D, 1:100), FoxP3-Spark 685 (206D, BioLegend, 1:200), and Bcl6-R718 (K112-91, BD Horizon, 1:200) for 1h at 25°C, and washed twice with True Nuclear Permeabilization/Wash buffer. Samples were resuspended in P2 and acquired on an Aurora using SpectroFlo v2.2 (Cytek). Flow cytometry data were analyzed using FlowJo v10 (Treestar).

For sorting PB, PBMC collected 1 week post-boost were incubated for 30 min on ice with purified CD16 (3G8, BioLegend, 1:100), CD32 (FUN-2, BioLegend, 1:100), and CD64 (10.1, BioLegend, 1:100), then stained for 30 min on ice with CD4-Spark UV 387 (SK3, 1:200), CD20-Pacific Blue (2H7, 1:400), CD71-FITC (CY1G4, 1:200), IgD-PerCP-Cy5.5 (IA6-2, 1:200), CD19-PE (HIB19, 1:200), CXCR5-PE-Dazzle 594 (J252D4, 1:50), CD38-PE-Fire 810 (HIT2, 1:200), CD14-A700 (HCD14, 1:200), and Zombie NIR (all BioLegend) diluted in P2. Cells were washed twice, and PB (live singlet CD4^−^ CD14^−^ CD19^+^ IgD^lo^ CD20^lo^ CD38^+^ CXCR5^lo^ CD71^+^) were sorted using a Bigfoot (Invitrogen) into Buffer RLT Plus (Qiagen) supplemented with 143 mM β- mercaptoethanol (Sigma-Aldrich) and immediately frozen on dry ice.

For sorting memory B cells, PBMC collected at baseline and 17 weeks after boosting were incubated for 30 min on ice with purified CD16 (3G8, BioLegend, 1:100), CD32 (FUN-2, BioLegend, 1:100), and CD64 (10.1, BioLegend, 1:100), then stained for 30 min on ice with CD4- Spark UV 387 (SK3, 1:200), IgD-BV421 (IA6-2, 1:200), CD3-FITC (HIT3a, 1:200), CD19-PE (HIB19, 1:200), CD27-PE-Fire 810 (O323, 1:200), CD14-A700 (HCD14, 1:200), and Zombie NIR (all BioLegend) diluted in P2. Cells were washed twice, and MBC (live singlet CD3^−^ CD4^−^ CD14^−^ CD19^+^ IgD^lo^) were sorted using a Bigfoot (Invitrogen) into Buffer RLT Plus (Qiagen) supplemented with 143 mM β-mercaptoethanol (Sigma-Aldrich) and immediately frozen on dry ice.

For bulk sorting GC B cells and LNPCs, lymph node FNA samples collected 8 weeks post- boosting were incubated for 30 min on ice with purified CD16 (3G8, BioLegend, 1:100), CD32 (FUN-2, BioLegend, 1:100), and CD64 (10.1, BioLegend, 1:100) in P2, then stained for 30 min on ice with PD-1-BB515 (EH12.1, BD Horizon, 1:100), CD20-Pacific Blue (2H7, 1:400), CD19- BV750 (HIB19, 1:100), IgD-PerCP-Cy5.5 (IA6-2, 1:200), CD71-PE (CY1G4, 1:400), CXCR5-PE-Dazzle 594 (J252D4, 1:50), CD38-PE-Cy7 (HIT2, 1:200), CD4-A700 (SK3, 1:400), and Zombie Aqua (all BioLegend) diluted in Brilliant Staining Buffer (BD Horizon). Cells were washed twice, and total GC B cells (live singlet CD4^−^ CD19^+^ IgD^lo^ CD20^+^ CD38^int^ CXCR5^+^ CD71^+^) and LNPCs (live singlet CD4^−^ CD19^+^ IgD^lo^ CD20^lo^ CD38^+^ CXCR5^lo^ CD71^+^) were sorted using a Bigfoot (Invitrogen) into Buffer RLT Plus (Qiagen) supplemented with 143 mM β- mercaptoethanol (Sigma-Aldrich) and immediately frozen on dry ice.

### Single-cell RNA-seq library preparation and sequencing

LN FNA samples were processed using the following 10x Genomics kits: Chromium Next GEM Single Cell 5′ Kit v2 (PN-1000263); Chromium Next GEM Chip K Single Cell Kit (PN-1000286); BCR Amplification Kit (PN-1000253); Dual Index Kit TT Set A (PN-1000215). Chromium Single Cell 5′ Gene Expression Dual Index libraries and Chromium Single Cell V(D)J Dual Index libraries were prepared according to manufacturer’s instructions. Both gene expression and V(D)J libraries were sequenced on a Novaseq S4 (Illumina), targeting a median sequencing depth of 50,000 and 5,000 read pairs per cell, respectively.

### Bulk BCR library preparation and sequencing

RNA was purified from sorted IgD^lo^ PBMCs (MBC), PB, GC B cells, LNPCs using the RNeasy Plus Micro kit (Qiagen). Reverse transcription, unique molecular identifier (UMI) barcoding, cDNA amplification (New England Biolabs #E6421), and Illumina linker addition to B cell heavy chain transcripts were performed using the human heavy chain only primers of NEBNext Immune Sequencing Kit (New England Biolabs #E6320) according to the manufacturer’s instructions (provided upon request). High-throughput 2x300bp paired-end sequencing was performed on the Illumina MiSeq platform with a 30% PhiX spike-in according to manufacturer’s recommendations, except for performing 325 cycles for read 1 and 275 cycles for read 2.

### Preprocessing of bulk sequencing BCR reads

Preprocessing of demultiplexed pair-end reads was performed using pRESTO v.0.6.2 (*67*) as previously described (*68*), with the exception that sequencing errors were corrected using the UMIs as they were without additional clustering (**Supplementary Data Table S2**).

### Preprocessing of 10x Genomics single-cell BCR reads

Demultiplexed pair-end FASTQ reads were preprocessed using Cell Ranger v.6.0.1 as previously described (*69*) (**Supplementary Data Table S3**).

### V(D)J gene annotation and genotyping

Initial germline V(D)J gene annotation was performed on the preprocessed BCRs using IgBLAST v.1.17.1 (*70*) with the deduplicated version of IMGT/V-QUEST reference directory release 202113-2 (*71*). Isotype annotation for 10x Genomics sequences was pulled from the ‘c_call’ column in the ‘filtered_contig_annotations.csv’ files outputted by Cell Ranger. Further sequence-level and cell-level quality controls were performed as previously described (*69*). Individualized genotypes were inferred based on sequences that passed all quality controls using TIgGER v.1.0.0 (*72*) and used to finalize V(D)J annotations. Sequences annotated as non-productively rearranged by IgBLAST were removed from further analysis.

### Clonal lineage inference

B cell clonal lineages were inferred on a by-individual basis based on productively rearranged sequences as previously described (*69*). Briefly, heavy chain-based clonal inference (*73*) was performed by partitioning the heavy chains of bulk and single-cell BCRs based on common V and J gene annotations and CDR3 lengths, and clustering the sequences within each partition hierarchically with single linkage based on their CDR3s (*74*). Sequences within 0.15 normalized Hamming distance from each other were clustered as clones. Following clonal inference, full-length clonal consensus germline sequences were reconstructed using Change-O v.1.0.2 (*75*). Within each clone, duplicate IMGT-aligned V(D)J sequences from bulk sequencing were collapsed using Alakazam v1.1.0 (*75*) except for duplicates derived from different time points, tissues, B cell compartments, isotypes, or biological replicates.

### BCR analysis

For B cell compartment labels, gene expression-based cluster annotation was used for single-cell BCRs; and FACS-based sorting and magnetic enrichment were used for bulk BCRs, except that IgD^lo^ enriched B cells from PMBC were labelled MBCs. For analysis involving the memory compartment, the memory sequences were restricted to those from blood. A heavy chain-based B cell clone was considered S-specific if it contained any sequence corresponding to a recombinant monoclonal antibody that was synthesized based on the single-cell BCRs and that tested positive for S-binding. Somatic hypermutation (SHM) frequency was calculated for each heavy chain sequence using SHazaM v.1.0.2 (*75*) by counting the number of nucleotide mismatches from the germline sequence in the variable segment leading up to the CDR3.

### Processing of 10x Genomics single-cell 5′ gene expression data

Demultiplexed pair-end FASTQ reads were first preprocessed on a by-sample basis and samples were subsequently subsampled to the same effective sequencing length and aggregated using Cell Ranger v.6.0.1 as previously described (*69*). Quality control was performed on the aggregate gene expression matrix consisting of 99,484 cells and 36,601 features using SCANPY v.1.7.2 (*76*). Briefly, to remove presumably lysed cells, cells with mitochondrial content greater than 20% of all transcripts were removed. To remove likely doublets, cells with more than 8,000 features or 80,000 total UMIs were removed. To remove cells with no detectable expression of common endogenous genes, cells with no transcript for any of a list of 34 housekeeping genes (*69*) were removed. The feature matrix was subset, based on their biotypes, to protein-coding, immunoglobulin, and T cell receptor genes that were expressed in at least 0.05% of the cells in any sample. The resultant feature matrix contained 14,796 genes. Finally, cells with detectable expression of fewer than 200 genes were removed. After quality control, there were a total of 95,915 cells from 10 single-cell samples (**Supplementary Data Table S3**).

### Single-cell gene expression analysis

Transcriptomic data was analyzed using SCANPY v.1.7.2 (*76*) as previously described (*69*) with minor adjustments suitable for the current data. Briefly, overall clusters were first identified using Leiden graph-clustering with resolution 0.18 (**Supplementary Fig. S2A, Supplementary Data Table S4**). UMAPs were faceted by participant and inspected for convergence to assess whether there was a need for integration. Cluster identities were assigned by examining the expression of a set of marker genes (*77*) for different cell types (**Supplementary Fig. S2B**). To remove potential contamination by platelets, 143 cells with a log-normalized expression value of >2.5 for *PPBP* were removed. Cells from the overall B cell cluster were further clustered to identify B cell subsets using Leiden graph-clustering resolution 0.25 (**Fig. 2A, Supplementary Table S4**). Cluster identities were assigned by examining the expression of a set of marker genes (*77*) for different B cell subsets (**Supplementary Fig. S2C-D**) along with the availability of BCRs. A group of 813 cells displaying expression signatures of both naïve and memory B cells was further clustered into 370 naïve B cells and 443 MBCs. Despite being clustered with B cells during overall clustering, one group tended to have both BCRs and relatively high expression levels of *CD2* and *CD3E*. Within this group, 4 cells with *CD3E* expression below group mean and *RGS13* expression above group mean were assigned to be GC B cells, whereas the rest “B & T”. Two unassigned groups tended to have no BCRs and no distinct expression signature of known B cell subsets. The “B & T” and unassigned groups were excluded from the final B cell clustering. Heavy chain SHM frequency and isotype usage of the B cell subsets were inspected for consistency with expected values to further confirm their assigned identities.

### Selection of single-cell BCRs from GC B cell or LNPC clusters for expression

Single-cell gene expression analysis was performed on a by-participant basis. Clonal inference was performed based on paired heavy and light chains. From every clone containing a cell from the GC B cell cluster and/or the LNPC cluster, one GC B cell or LNPC was selected. For selection, where a clone spanned both the GC B cell and LNPC compartments, a compartment was first randomly selected. Within that clone, the cell with the highest heavy chain UMI count was then selected, breaking ties based on *IGHV* SHM frequency. In all selected cells, native pairing was preserved. The selected BCRs were curated as previously described prior to synthesis (*69*).

### Transfection for recombinant mAbs and Fab production

Selected pairs of heavy and light chain sequences were synthesized by GenScript and sequentially cloned into IgG1, Igκ/λ and Fab expression vectors. Heavy and light chain plasmids were co-transfected into Expi293F cells (Thermo Fisher Scientific) for recombinant mAb production, followed by purification with protein A agarose resin (GoldBio). Expi293F cells were cultured in Expi293 Expression Medium (Gibco) according to the manufacturer’s protocol.

### Chimeric VSV-SARS-CoV-2 neutralization assay

The VSV-based neutralization assay was performed as described previously (*43*). The gene encoding spike of SARS-CoV-2 isolate WA1/2020 (with D614G mutation) was synthesized and replaced the native envelope glycoprotein of an infectious molecular clone of VSV, and resulting chimeric viruses expressing S protein from SARS-CoV-2 D614G was used for GFP reduction neutralization tests as previously described (*43, 64*). Briefly, 2x10^3^ PFU of VSV-SARS-CoV-2- S**Δ**21 was incubated for 1 h at 37°C with recombinant mAbs diluted to 10 µg/ml. Antibody-virus complexes were added to Vero E6 cells in 96-well plates and incubated at 37°C for 7.5 h. Cells were subsequently fixed in 2% formaldehyde (Electron Microscopy Sciences) containing 10 mg/mL Hoechst 33342 nuclear stain (Invitrogen) for 45 min at room temperature, when fixative was replaced with PBS. Images were acquired with an Cytation C10 automated microscope (BioTek) using the DAPI and GFP channels to visualize nuclei and infected cells (i.e., eGFP-positive cells), respectively (4X objective, 4 fields per well, covering the entire well). Images were analyzed using the Gen5 3.12 software’s Data Reduction tool (BioTek). GFP-positive cells were identified in the GFP channel following image preprocessing, deconvolution and subsequently counted within the Gen5 3.12 software. The sensitivity and accuracy of GFP-positive cell number determinations were validated using technical replicates. The percent infection reduction was calculated from wells to which no antibody was added. A background number of GFP-positive cells was subtracted from each well using an average value determined from at least 4 uninfected wells.

### Viruses

The WA1/2020 recombinant strain with D614G substitution was described previously (*78, 79*). The BA.1 isolate (hCoV-19/USA/WI-WSLH-221686/2021) was obtained from an individual in Wisconsin as a mid-turbinate nasal swab (*80*). BA.2.86 (hCoV-1/USA/MI-UM- 10052670540/2023), XBB.1.5(hCoV-19/USA/MD-HP40900-PIDYSWHNUB/2022), JN.1(hCoV-19/USA/CA-Stanford-165_S10/2023), HV.1 (hCoV-19/USA/CA-Stanford- 165_S45/2023), and KP.2 (hCoV-19/USA/CA-Stanford-181_S33/2024) were generous gifts from A. Pekosz (Johns Hopkins), and M. Suthar (Emory University).

The EG.5.1 variant of SARS-CoV-2 (hCoV-19/USA/CA-Stanford-147_S01/2023) was propagated on Vero-hTMPRSS2 cells. The virus stocks were subjected to next-generation sequencing, and the S protein sequences were identical to the original isolates. The infectious virus titer was determined by plaque and focus-forming assay on Vero-hACE2-hTMPRSS2 or Vero- hTMPRSS2 cells.

### Hamster challenge studies

Animal studies were carried out in accordance with the recommendations in the Guide for the Care and Use of Laboratory Animals of the National Institutes of Health. The protocols were approved by the Institutional Animal Care and Use Committee at the Washington University School of Medicine (assurance number A3381–01). Five-week-old male hamsters (n = 6/group) were obtained from Charles River Laboratories and housed in a biosafety level 3 facility at Washington University. One day prior to challenge with 10^4^ plaque forming units (PFU) of EG.5.1, the animals received intraperitoneal 10 mg/kg of mAb-52 or isotype control (1G05) in PBS. Animal weights were measured daily for the duration of the experiment. Three days after the challenge, hamsters were euthanized and necropsied, and the left lung lobe, nasal wash and nasal turbinate’s were collected for virological analysis. These tissues were homogenized in 1.0 mL of DMEM, clarified by centrifugation (1,000 × g for 5 min) and used for viral titer analysis by quantitative RT-PCR (RT-qPCR) using primers and probes targeting the N gene, and by plaque assay.

### Virus titer assays

Plaque assays were performed on Vero-hACE2-hTMPRSS2 cells in 24-well plates. Lung tissue and nasal turbinate homogenates were serially diluted 10-fold, starting at 1:10, in cell infection medium (DMEM supplemented with 2% FBS, 10 mM HEPES, and 2 mM L-glutamine). Two hundred and fifty microliters of the diluted homogenate were added to a single well per dilution per sample. After 1 h at 37 °C, the inoculum was aspirated, the cells were washed with PBS, and a 1% methylcellulose overlay in MEM supplemented with 2% FBS was added. Seventy-two hours after virus inoculation, the cells were fixed with 4% formalin, and the monolayer was stained with crystal violet (0.5% w/v in 25% methanol in water) for 1 h at 20 °C. The number of plaques were counted and used to calculate the PFU/mL. To quantify viral RNA levels in the homogenates, RNA was extracted from 100 µL homogenate, using an automated RNA extraction machine (KingFisher Flex) and the MagMax Viral Pathogen kit according to the manufacturer’s recommendations. The RNA was eluted in 50 µL of water. Four microliters RNA was used RT-qPCR to detect and quantify N gene of SARS-CoV-2 using the TaqMan RNA-to-CT 1-Step Kit (Thermo Fisher Scientific) with the following primers and probes for the N-gene, Forward primer: ATGCTGCAATCGTGCTACAA; Reverse primer: GACTGCCGCCTCTGCTC; Probe: /56-FAM/TCAAG GAAC/ ZEN/AACATTGCCAA/3IABkFQ/. Viral RNA was expressed as gene copy numbers per mg for lung tissue homogenates and per mL for nasal turbinates, based on a standard included in the assay (*50, 81–83*).

### Focus reduction neutralization test (FRNT)

Serial dilutions of each mAb were incubated with 10^2^ focus-forming units (FFU) of different SARS-CoV-2 strains (WA1/2020 D614G, BA.1, XBB.1.5, EG.5.1, BA.2.86, HV.1, JN.1, or KP.2) for 1 h at 37°C. Antibody-virus complexes were added to Vero-TMPRSS2 cell monolayers in 96- well plates and incubated at 37°C for 1 h. Subsequently, cells were overlaid with 1% (w/v) methylcellulose in MEM. Plates were harvested 30 h (WA1/2020 D614G) or 65-70 h (Omicron strains) later by removing overlays and fixed with 4% PFA in PBS for 20 min at room temperature. Plates were washed and incubated with an oligoclonal pool of anti-S antibodies (SARS2–2, SARS2–11, SARS2–16, SARS2–31, SARS2–38, SARS2–57, and SARS2–71), and an additional oligoclonal pool of anti-S antibodies with Supplementary reactivity (SARS2–08, −09, −10, −13, −14, −17, −20, −26, 27, −28, −31, −41, −42, −44, −49, −62, −64, −65, and −67) were included for staining plates infected with Omicron strains. Plates were subsequently incubated with HRP-conjugated goat anti-mouse IgG (Sigma Cat # A8924, RRID: AB_258426) in PBS supplemented with 0.1% saponin and 0.1% bovine serum albumin. SARS-CoV-2-infected cell foci were visualized using TrueBlue peroxidase substrate (KPL) and quantitated on an ImmunoSpot microanalyzer (Cellular Technologies).

### Antibody escape mapping using deep mutational scanning

XBB.1.5 RBD deep mutational scanning libraries were designed as described previously (*54*). For antibody selection experiments, approximately 1 M transcription units of the library were incubated with 5.3 and 21.3 µg/ml of mAb-52 for 45 min at 37C. These concentrations were determined using XBB.1.5 pseudovirus neutralization assay (*54*) and were approximately IC99*4 and IC99*16 as measured on HEK-293T-ACE2 cells. After incubation virus-antibody mix was used to infect HEK-293T-ACE2 cells. Viral genomes were recovered for deep sequencing 12 hours after infection. Antibody escape was mapped using two independent XBB.1.5 RBD libraries. Mutation-level escape was determined by using non-neutralizable control as described previously (*84*) and a biophysical model implemented in *polyclonal* (*85*) package https://jbloomlab.github.io/polyclonal/.

Interactive escape plots for mAb-52 can be found at https://dms-vep.org/SARS-CoV-2_XBB.1.5_RBD_DMS_mAB-52/htmls/mAb_52_mut_effect.html and the full analysis pipeline used to map escape can be found at https://github.com/dms-vep/SARS-CoV-2_XBB.1.5_RBD_DMS_mAB-52.

### Fab generation

Fabs of mAb-52 used for binding studies were produced in-house as described previously (*43*). Heavy chain encoding plasmids were restriction digested with *AgeI*, *SalI* and VH region was gel extracted and subcloned by ligation into Fab expression vector. Genes encoding germline revertants of FAb-52 were synthesized by Genscript and subcloned into heavy chain encoding plasmids as described above. The sequence confirmed Fab heavy chain and light chain plasmids were co-transfected into Expi293F cells (Gibco) for expression and purified with HisPur Ni-NTA resin (Thermo Scientific).

### Biolayer interferometry

Kinetic binding studies were performed on an Octet-R8 (Sartorius) instrument. Avi-tagged biotinylated RBD of SARS-CoV-2 variants WA1/2020, BA.1, EG.5.1, HV.1, JN.1, KP.2, KP.3 were produced inhouse. Octet ™ SA-Biosensor tips (Sartorius) were pre-equilibrated in HBS supplemented with 0.05% Tween-20 and 1% BSA (kinetic buffer A) followed by loading of Avi- RBD proteins to 1.0 nm. Kinetic binding studies were performed in kinetic buffer A by monitoring Fabs association (200 s) and dissociation (600 s). Octet ™ SA-Biosensors that were not loaded were used as reference sensor. Kinetic parameters of reference subtracted kinetic traces were calculated with Octet BLI analysis software v12.1 using a global fit 1:1 binding model. Traces were plotted with GraphPad Prism v10.

BLI competition studies were performed on an Octet Red instrument (ForteBio). Biotinylated Avi tagged WA1 RBD was loaded onto Streptavidin sensor tips (Sartorius) to 2 nm that were pre-equilibrated in kinetic buffer A. Following loading, mAb-52/1G05 (isotype) (250 nM) were monitored for binding for 200 s and followed by 200 s of competitive binding against 2B04 (class I/A) and ACE2 receptor (250 nM). Octet™ SA-Biosensors that were loaded and dipped in blank buffer were used as reference sensors. The relative shift in competitive mAb/receptor binding was quantified between 200-400 s relative to 1G05 isotype control following reference subtraction of kinetic traces. Traces were plotted with GraphPad Prism v10.

### Cryo electron microscopy sample preparation, data collection

The SARS-CoV-2 XBB.1.5 HexaPro spike (GISAID Accession ID-EPI_ISL_18416647) was mixed with Fab-52 at a concentration of 2 mg/mL, using a 1.5 molar excess of Fab, and incubated for 20 minutes at room temperature. Immediately before grid preparation, fluorinated octyl-maltoside was added to the complex at a final concentration of 0.02% wt/vol. Next, 3 μl aliquots were applied to UltrAuFoil gold R1.2/1.3 grids, which were blotted for 6 seconds at a blot force of 0, at 22°C and 95% humidity. The samples were then plunge-frozen in liquid ethane using a Vitrobot Mark IV system (ThermoFisher Scientific). Imaging was conducted on a Titan Krios microscope operated at 300 kV and equipped with a 15 eV energy filter and a Gatan K3 direct electron detector. A total of 6,614 movie frames were captured, with a cumulative dose of 48.95 e-/Å²/s. Images were recorded at a magnification of 105,000, corresponding to a calibrated pixel size of 0.4125 Å/pixel, with a defocus range from -0.9 to -2.3 μm.

### Cryo electron microscopy data processing, structure modelling and refinement

The movies were aligned and dose-weighted using the patch motion correction feature in cryoSPARC v4.3.1 (*86*). The contrast transfer function (CTF) was estimated using Patch CTF, and particles were picked with cryoSPARC’s template picker. The picked particles were extracted with a box size of 1024 pixels, with 4× binning, and subjected to a 2D classification. An initial model was generated from 625,012 selected particles, and the best class containing 267,722 was chosen for further analysis. After two rounds of non-uniform refinement, without imposed symmetry, the particles were subjected to 3D classification with six classes, where one class was selected for additional processing, containing 317,638 particles. These particles were re-extracted with a box size of 1024 pixels and 2× binning, followed by further rounds of non-uniform refinement that included local and global CTF refinement, resulting in a final global map with a nominal resolution of 2.58 Å. There was only one Fab with one of the protomers that was subjected to local refinement with a soft mask extended by 6 pixels and padded by 12 pixels encompassing the receptor binding domain (RBD) and Fab. This local refinement yielded a resolution of 3.11 Å. The two half-maps from this refinement were sharpened using DeepEMhancer (*87*). The reported resolutions are based on the gold-standard Fourier shell correlation criterion of 0.143.

The focused maps sharpened with DeepEMhancer were used for model building. The initial model was created using ModelAngelo (*88*) and then manually refined with COOT (*89*). N-linked glycans were added manually in COOT using the glyco extension. The model underwent further refinement in Phenix, employing real-space refinement, and was validated using MolProbity (*90*) (**Supplementary Data Table S5**). The structural biology software was compiled and made available through SBGrid (*91*).

### Quantification and statistical analysis

Statistics were employed in the manuscript as described in the Figure legends.

## Acknowledgements

We thank the generous participation of the donors for providing specimens. We thank Lisa Kessels and the Washington University School of Medicine 382 Study Team (study coordinators Alem Haile, Ryley Thompson, Delaney Carani, RN, Kim Gray, MSN, APRN-BC, and Chapelle Ayres; pharmacists Michael Royal, RPh and John Tran; and laboratory technicians Laura Blair, Anita Afghanzada, and Natalie Schodl) for assistance with scheduling participants and sample collection. We thank Pamela Woodard, Betsy Thomas, Mike Harrod, Rosemary Hamlin, Maggie Rohn, and the staff of the Center for Clinical Research Imaging at Washington University School of Medicine for assistance with sample collection. We thank Claire Dalton and Brittany Roemmich for performing the nucleocapsid binding assay. We also acknowledge support from the Irma T. Hirschl/Monique Weill-Caulier Trust. We thank support of the computational and data resources and staff expertise provided by Scientific Computing and Data at the Icahn School of Medicine at Mount Sinai. We thank Hanover Matz for critically reading the manuscript. We also thank all the members of the Ellebedy lab for their valuable suggestions. The WU382 study was reviewed and approved by the Washington University Institutional Review Board (approval no. 202109021).

## Funding and resources

National Institutes of Health grant P01AI172531 (AHE)

National Institutes of Health grant 75N93021C00014 Option 22A (AHE) National Institutes of Health grant R01AI168178 (AHE)

National Institutes of Health grant P01AI168347 (AHE)

National Institutes of Health grant R01 AI168178 (GB and AHE).

National Institutes of Health, Collaborative Influenza Vaccine Innovation Centers contract 75N93019C00051 (GB).

NYU Langone Health’s Cryo-Electron Microscopy Laboratory, RRID SCR_019202.

National Institutes of Health, Laura and Isaac Perlmutter Cancer Center Support Grant NIH/NCI P30CA016087.

National Center for Advancing Translational Sciences, Clinical and Translational Science Awards grant UL1TR004419 (GB).

National Institutes of Health, Office of Research Infrastructure award S10OD026880 and S10OD030463.

National Institutes of Health, Center of Excellence for Influenza Research and Response (CEIRR) contract 75N93021C00016 (ACMB, MSD, AHE).

## Author contributions

Conceptualization: AHE, JAO, RMP, RP, BN, and SC Data curation: SKM, JQZ, and JST

Formal analysis: AHE, GB, MSD, ACMB, JDB, SKM, DJ, BY, TLD, BD, JQZ, and JST,

Funding acquisition: AHE, GB, MSD, ACMB, and JDB

Investigation: SKM, DJ, BY, WBA, TLD, BD, AC, SCH, JQZ, WK, JST, AJS, and FH Methodology: AHE, GB, ACMB, MSD, JDB, RMP, MKK, JAO, SKM, JST, AJS, and JQZ.

Project administration: AHE, RMP, JAO, and MKK

Resources: AHE, GB, MSD, JDB, ACMB, RMP, JAO, MKK, RN, DKE, BSS, WDM, SMS, and CWF

Supervision: AHE, GB, MSD, and JDB Validation: SKM, DJ, BY, TLD, BD, and JQZ Visualization: SKM, WBA, DJ, BD, and JQZ Writing – original draft: SKM, and AHE

Writing – review & editing: SKM, DJ, BY, WBA, TLD, BD, AC, SCH, JQZ, WK, JST, AJS, FH, SMS, CWF, RN, BN, SC, MKK, DKE, RP, BSS, WDM, JAO, RMP, JDB, ACMB, MSD, GB, and AHE

## Data and material availability

Upon acceptance, raw sequencing data and transcriptomics count matrix will be deposited at Sequence Read Archive and Gene Expression Omnibus under BioProject xxxxx. Processed BCR and transcriptomics data will be deposited at Zenodo (https://doi.org/10.5281/zenodo.xxxxx). Materials are available upon request, through a simple interinstitutional materials transfer agreement. The EM maps have been deposited in the Electron Microscopy Data Bank (EMDB) under accession code EMD-47426 and the accompanying atomic coordinates in the Protein Data Bank (PDB) under accession code 9E21. The aligned micrographs are available on the Electron Microscopy Public Image Archive (EMPIAR) under accession number EMPIAR-12414. The content is solely the responsibility of the authors and does not necessarily represent the official views of the National Institutes of Health.

## Supplementary Materials

Figs. S1 to S10 Tables S1 to S5

