## Supplementary Fig. S1, S2, S3, S4, S5, S6, S7, S8, S9, S10, Supplementary Data Table S1, S2, S3,S4, S5 for "Defining a highly conserved B cell epitope in the receptor binding motif of SARS-CoV-2 spike glycoprotein"

### Supplementary Figures

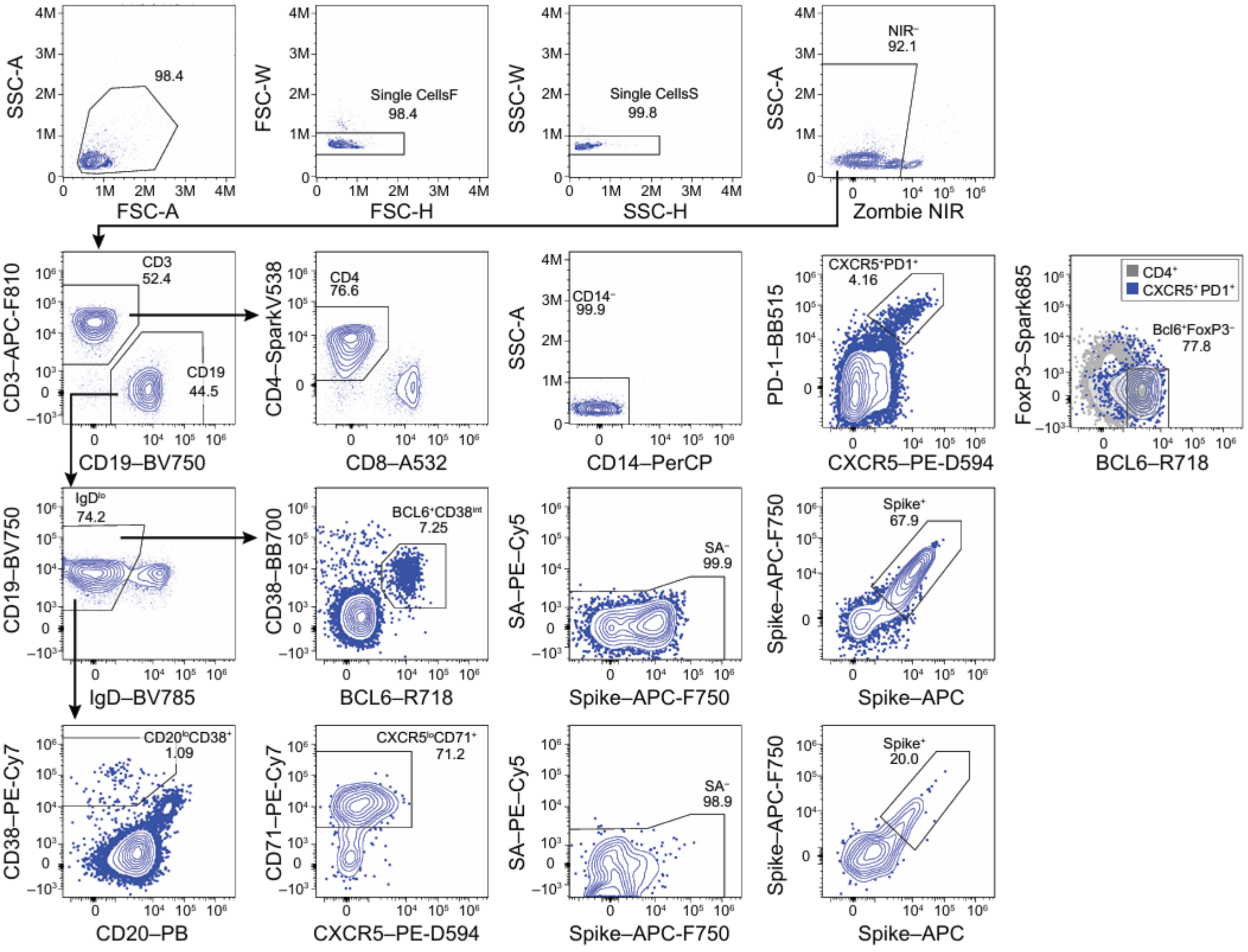


**Supplementary Figure S1. Lymph node fine needle aspirate flow cytometry gating strategy.**

Representative flow plots to analyze $T_{FH}$, $S^+$ germinal centre B cells and lymph node plasma cells.

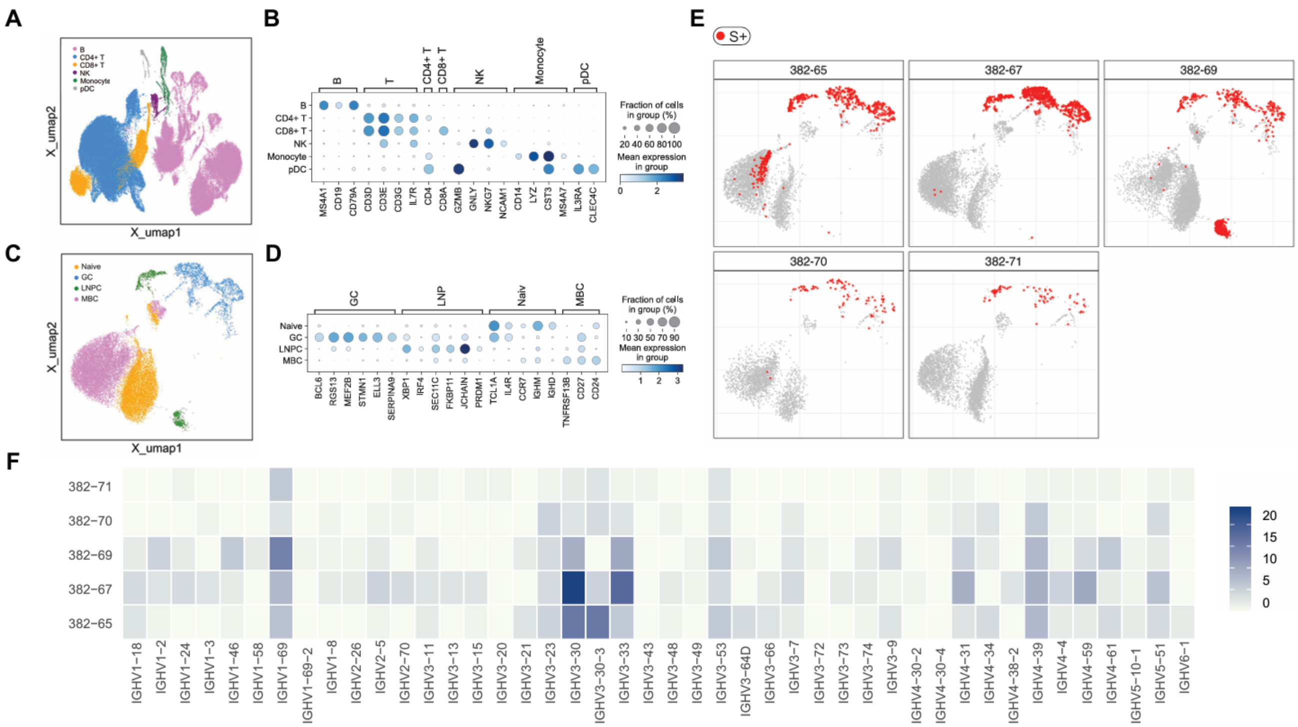

**Supplementary Figure S2. Lymph node fine needle aspirate scRNA-seq transcriptional clusters. (A, C)** UMAPs depicting entire lymph node transcriptional clusters **(A)** and B cells alone **(C)**. B cells found in S$^+$ clones in each participant are overlayed in red **(E)**. The marker genes used for annotating the transcriptional clusters are depicted as dot plots **(B, D)**. IGHV gene usage by S$^+$ B cell clones is visualized as a heatmap colored based on clone counts **(F)**.

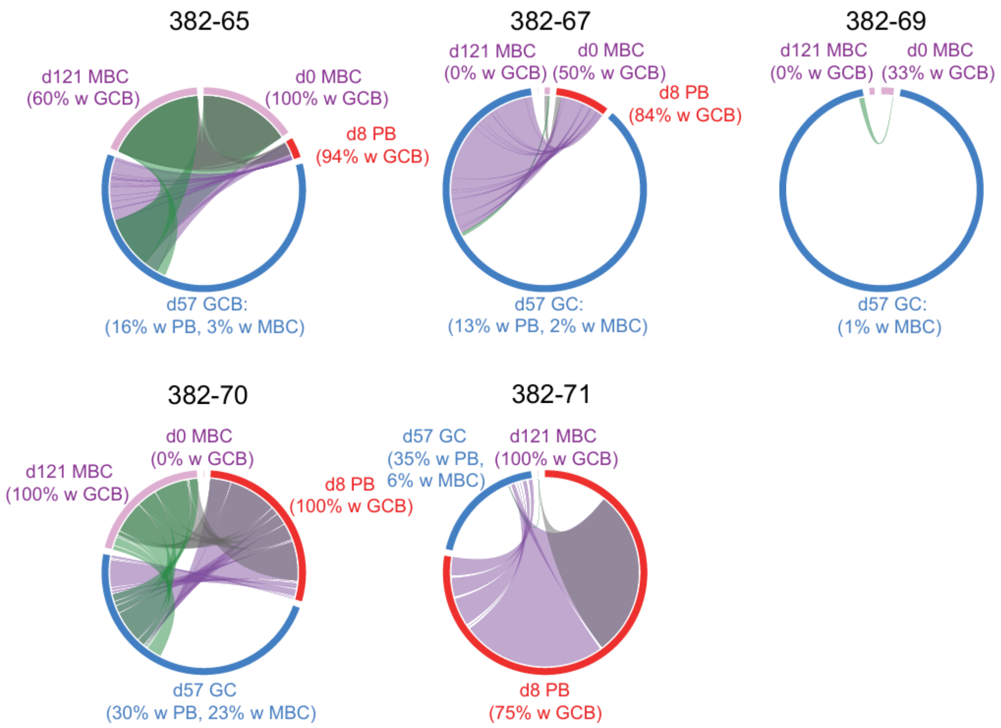


**Supplementary Figure S3. Longitudinal tracking of S-binding B cell clones across lymph node and peripheral blood compartments.** Circos diagrams showing clonal overlap between S-binding plasmablasts (PB, red arc), memory B cells (MBC, pink arc), and GC B cells (GCB, blue arc) compartment at indicated time points. Arcs represent B cell compartments, with arc lengths being proportional to the total number of B cells in each compartment. Chords represent clonal connections spanning at least two compartments, with the widths of chord ends corresponding to the number of B cells from the respective compartments. Purple, green and grey chords correspond to clonal connections spanning GCB-PB, GCB-MBC and PB-MBC compartments respectively. Percentages are of B cell clones in the compartment represented by the arc that were related to B cells in indicated compartment and time point.

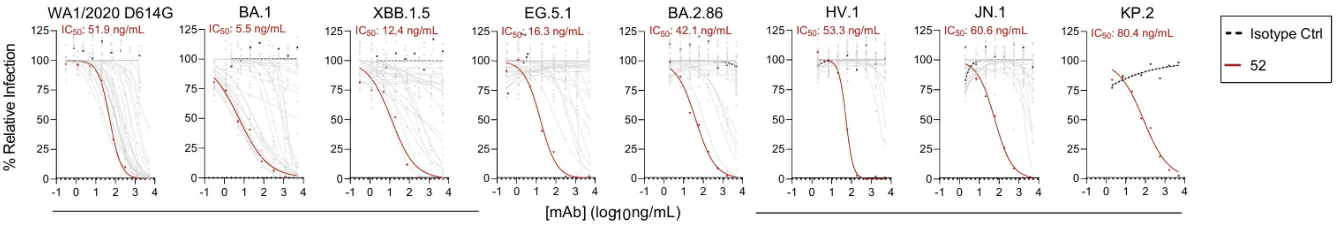


**Supplementary Figure S4. Authentic SARS-CoV-2 variant neutralizations.** Neutralization of mAbs against (**A**) WA1/2020 D614G, (**B**) BA.1, (**C**) XBB.1.5, (**D**) EG.5.1, (**E**) BA.2.86, (**F**) HV.1, (**G**) JN.1, and (**H**) KP.2. $IC_{50}$ values indicated above the graphs. Results are from technical duplicates of one experiment.

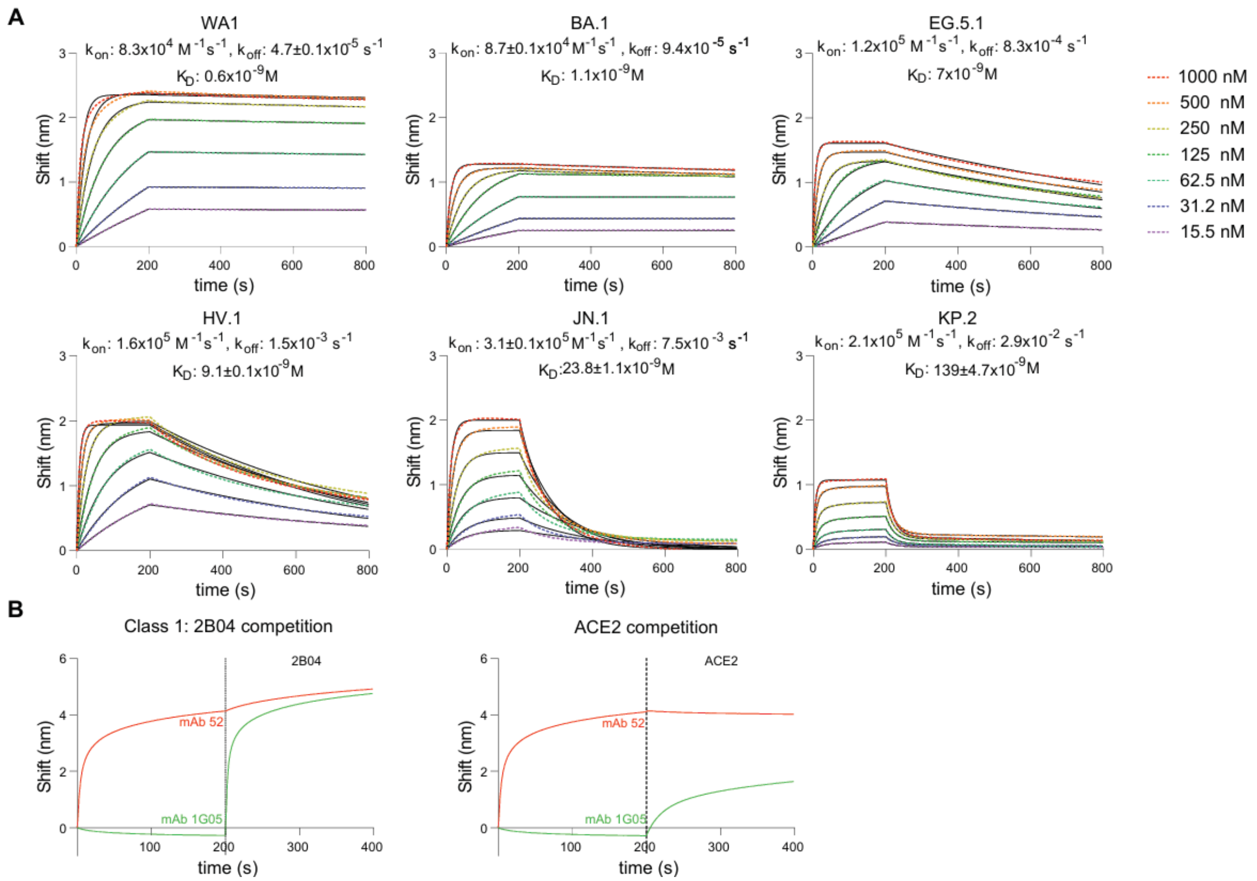


**Supplementary Figure S5. BLI binding affinity and competition of neutralizing antibodies. (A)** BLI binding affinity of mAb-52 binding receptor binding domain of WA1, BA.1, EG.5.1, HV.1, JN.1, KP.2. **(B)** BLI binding competition against class I/A-binding mAb 2B04 and ACE2. Results are from kinetic measurements of dilution series of one experiment.

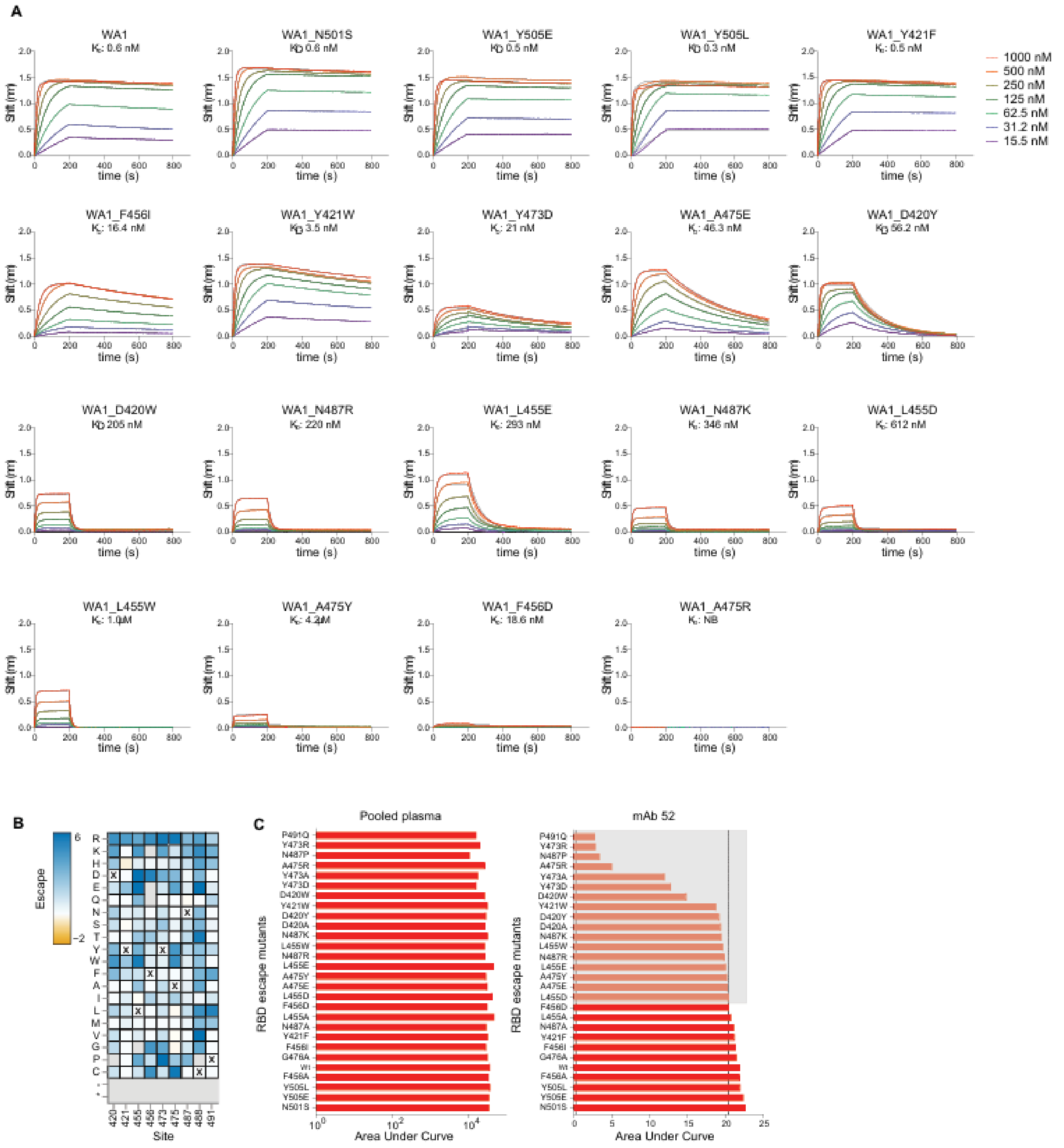


**Supplementary Figure S6. Escape mutant binding analysis. (A)** Fab-52 binding affinity against key RBD escape mutants identified in deep mutational scanning. **(B)** Key residue mutational escape heatmap in XBB.1.5 RBD library. XBB.1.5 wild type amino acid are depicted with an X mark, and amino acids in grey are absent in the library or highly deleterious. The complete data for line plot, heat map and codes can be accessed at dms-vep.org/SARS-CoV-2_XBB.1.5_RBD_DMS_mAB-52/htmls/mAb_52_mut_icXX.html **(C)** ELISA binding area under the curve (AUC) of mAb-52 and pooled plasma (d28) binding RBD escape mutants. RBD mutants binding mAb-52 with >10% reduction in overall binding were considered epitopic residues of mAb-52. The mutations highlighted in gray box are confirmed escape mutants. Results are from technical duplicates of one experiment.

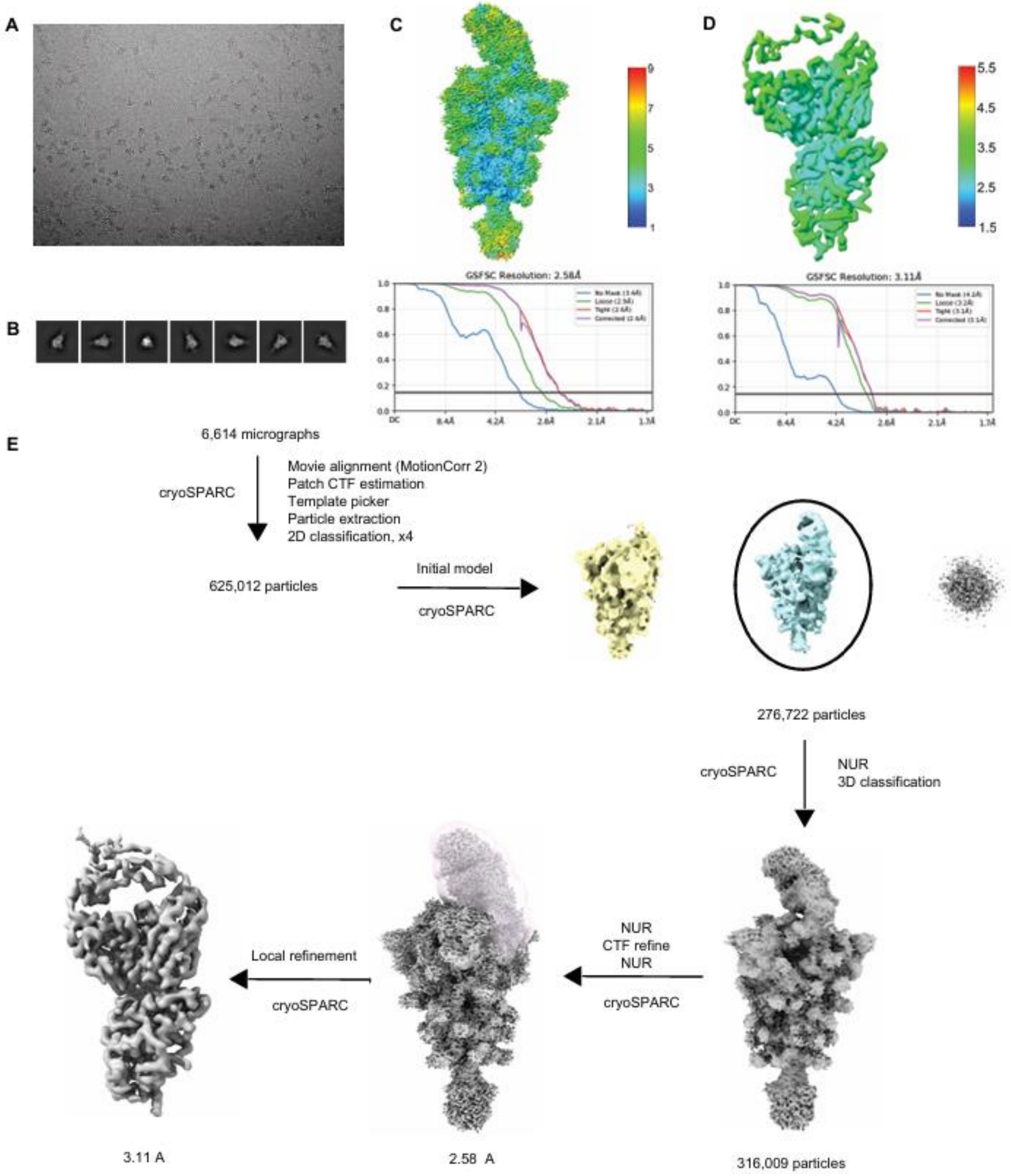


**Supplementary Figure S7. Overview of cryo-EM data processing and local resolution assessment. (A)** Representative electron micrograph. **(B)** 2D class average obtained for SARS-CoV-2 XBB.1.5 spike ectodomain in complex with 52 Fab. Local resolution and gold-standard

Fourier shell correlation curves generated with cryoSPARC v4.3.1 **(C)** for overall **(D)** and locally refined, RBD-Fab complex. **(E)** An outline of cryo-EM data processing workflow.

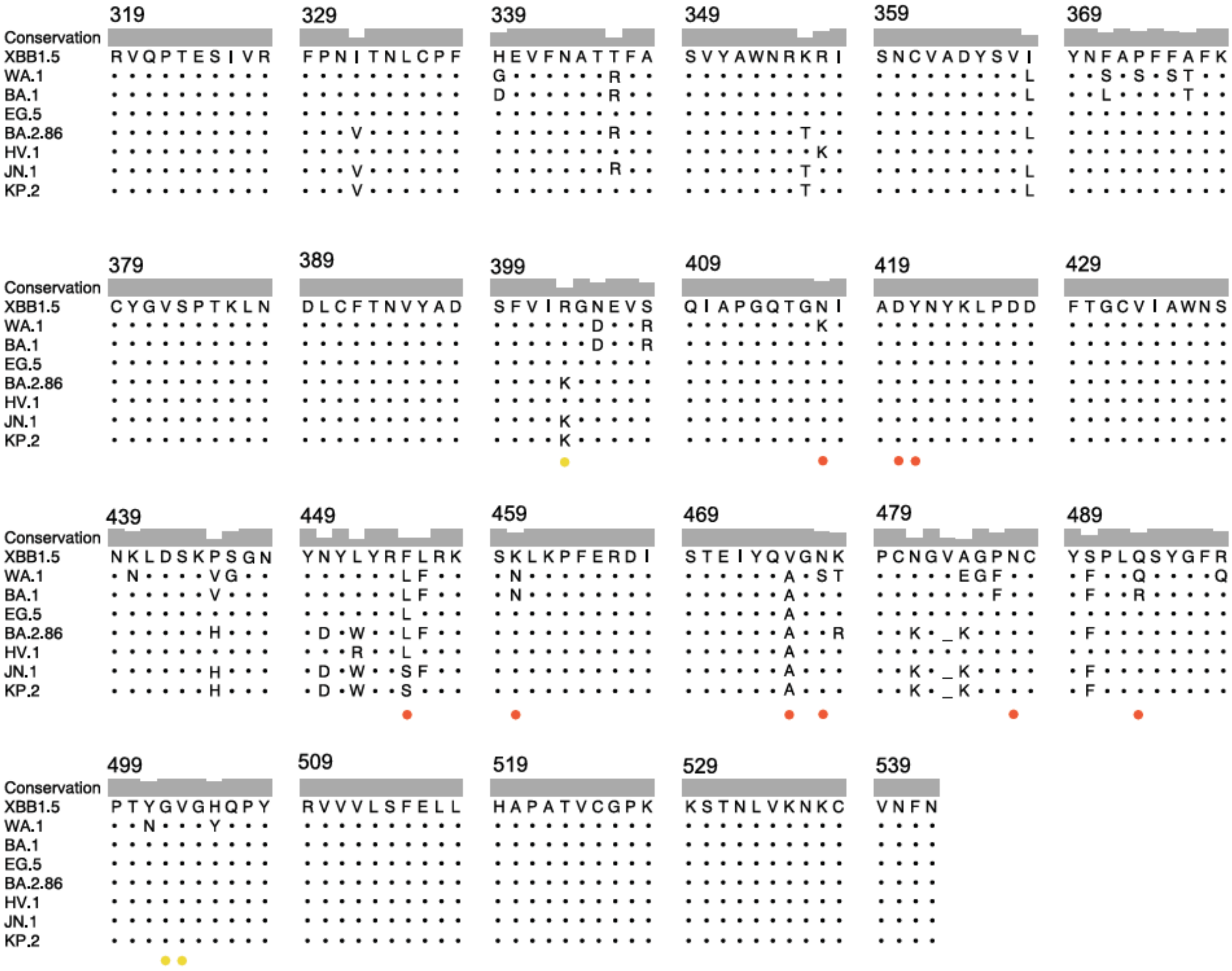

**Supplementary Figure S8. Multiple Sequence alignment**. SARS-CoV-2 spike RBD (aa 319-542) of XBB.1.5, WA.1, BA.1, EG.5, BA.2.86, HV.1, JN.1and KP.2. Epitope residues of Fab-52 are indicated with colored dots, orange red for heavy chain and yellow for light chain.

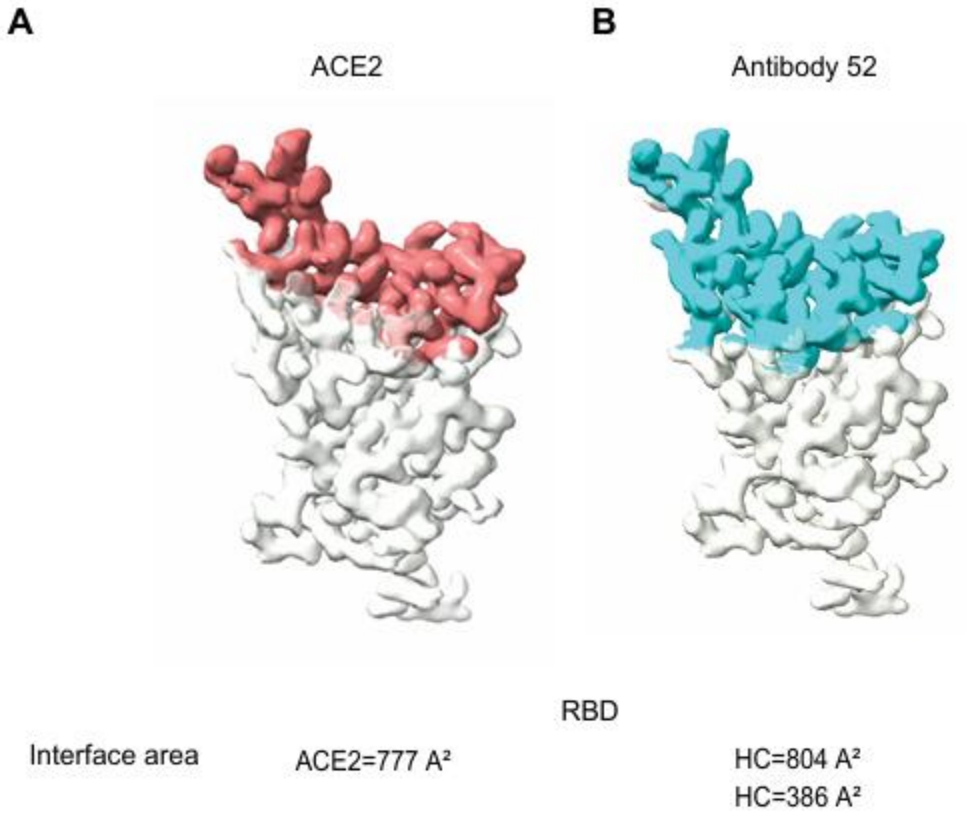


**Supplementary Figure S9. Buried surface area epitope surface representation. (A)** ACE2 (PDB-8IOU) and **(B)** mAb-52.

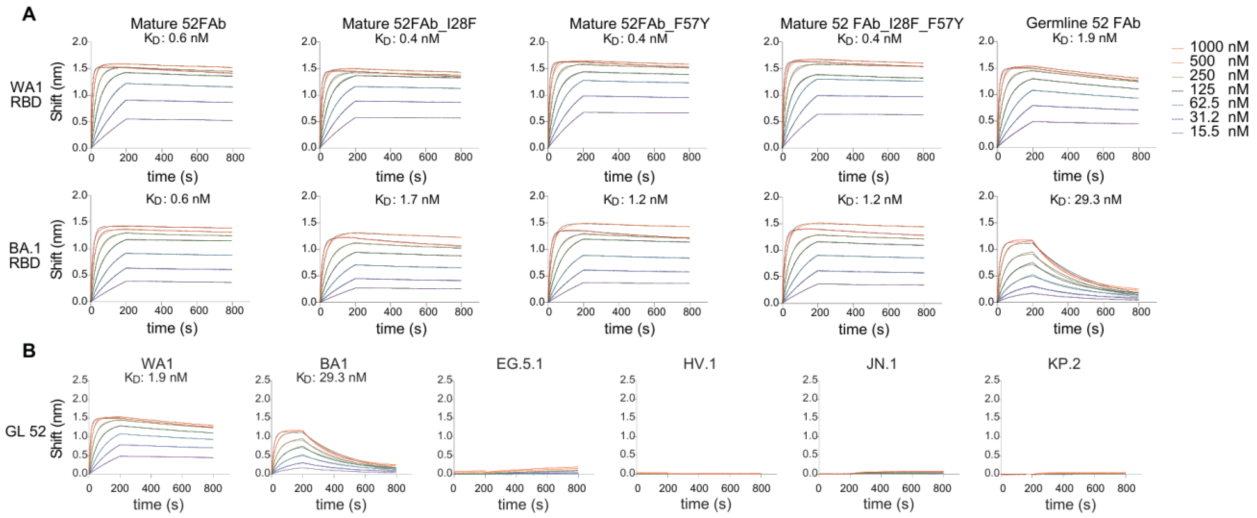


**Supplementary Figure S10. BLI binding affinity (A)** Mature, germline revertant and germline Fab-52 binding RBD of WA1 and BA.1. **(B)** Germline Fab-52 binding variant RBD of WA1, BA.1, EG.5.1, HV.1, JN.1, KP.2. Results are from kinetic measurements of dilution series of one experiment.

### Supplementary data tables

#### Supplementary Data Table S1. Study WU382 participant demographics.

Proportion of participants in indicated demographic categories.

| Variable | Total n = 9<br>n (%) |
|---|---|
| **Age (median [range])** | 51 (27-72) |
| **Sex** | |
| Female | 5 (55.6) |
| Male | 4 (44.4) |

**Supplementary data table S2. Processing of BCR reads from bulk sequencing**

| | | | | | Sequence Count | | | |
|---|---|---|---|---|---|---|---|---|
| Participant | Timepoint | Tissue | Sorting | Cell Count | Input Reads | Preprocessed Reads | Post-QC Productive Heavy Chains | Unique Heavy Chain VDJs |
| 382-65 | d0 | Blood | IgDlo | 250464 | 1474888 | 9670 | 8003 | 5268 |
| 382-65 | d8 | Blood | PB | 25078 | 1629948 | 10858 | 8952 | 4265 |
| 382-65 | d57 | LN | GC B | 28569 | 995015 | 5451 | 4295 | 2563 |
| 382-65 | d57 | LN | LNPC | 4146 | 1309457 | 15396 | 12329 | 4266 |
| 382-65 | d121 | Blood | IgDlo | 494579 | 1972488 | 17182 | 14866 | 9459 |
| 382-67 | d0 | Blood | IgDlo | 102255 | 1857265 | 12973 | 11231 | 7022 |
| 382-67 | d8 | Blood | PB | 21585 | 1733950 | 6762 | 5446 | 3506 |
| 382-67 | d57 | LN | GC B | 23203 | 949129 | 1995 | 1512 | 891 |
| 382-67 | d57 | LN | LNPC | 3266 | 1295174 | 6675 | 5160 | 2217 |
| 382-67 | d121 | Blood | IgDlo | 219064 | 1752127 | 11871 | 10300 | 6799 |
| 382-69 | d0 | Blood | IgDlo | 104705 | 1586882 | 12487 | 10657 | 6899 |
| 382-69 | d57 | LN | GC B | 1604 | 1052477 | 283 | 154 | 92 |
| 382-69 | d57 | LN | LNPC | 302 | 1053350 | 966 | 649 | 299 |
| 382-69 | d121 | Blood | IgDlo | 85444 | 1985842 | 16447 | 14578 | 9126 |
| 382-70 | d0 | Blood | IgDlo | 230258 | 2018408 | 19335 | 17377 | 9963 |
| 382-70 | d8 | Blood | PB | 59615 | 1945779 | 27737 | 25634 | 14410 |
| 382-70 | d57 | LN | GC B | 4801 | 1001226 | 996 | 649 | 341 |
| 382-70 | d57 | LN | LNPC | 1644 | 954374 | 4563 | 3567 | 1481 |
| 382-70 | d121 | Blood | IgDlo | 272610 | 2156732 | 64838 | 58829 | 23979 |
| 382-71 | d0 | Blood | IgDlo | 295954 | 1944729 | 17354 | 15430 | 10148 |
| 382-71 | d8 | Blood | PB | 51971 | 1852169 | 43305 | 39101 | 12593 |
| 382-71 | d121 | Blood | IgDlo | 300504 | 1736137 | 18442 | 16869 | 11592 |

**Supplementary data table S3. Processing of BCR and 5' gene expression data from scRNA-seq**

| Participant | Timepoint | Tissue | Replicate | BCR | | 5' gene expression | | | |
|---|---|---|---|---|---|---|---|---|---|
| | | | | Pre-QC number of cells | Post-QC number of cells | Pre-QC number of cells | Post-QC number of cells | Median number of UMIs per cell | Median number of genes per cell |
| 382-65 | d57 | LN | 1 | 2990 | 2750 | 9648 | 9082 | 3646 | 1419.5 |
| | d57 | LN | 2 | 2548 | 2350 | 8465 | 7907 | 3740 | 1410 |
| 382-67 | d57 | LN | 1 | 3266 | 2938 | 7360 | 6995 | 3536 | 1447 |
| | d57 | LN | 2 | 3710 | 3293 | 7409 | 7086 | 3382.5 | 1395 |
| 382-69 | d57 | LN | 1 | 4638 | 4205 | 9358 | 8680 | 3474 | 1404 |
| | d57 | LN | 2 | 8341 | 6494 | 22007 | 21591 | 776 | 514 |
| 382-70 | d57 | LN | 1 | 1508 | 1400 | 8982 | 8808 | 3228 | 1244 |
| | d57 | LN | 2 | 1399 | 1330 | 8733 | 8513 | 3392 | 1312 |
| 382-71 | d57 | LN | 1 | 2113 | 2018 | 8692 | 8563 | 3732 | 1444 |
| | d57 | LN | 2 | 2091 | 2002 | 8830 | 8690 | 3806.5 | 1456 |

**Supplementary data table S4. scRNA-seq derived transcriptional cluster cell counts and frequencies**

| Participant | Overall cluster | Cell count (% of total cells) | B cell cluster | Cell count (% of B cells) | SARS-CoV-2 S-binding cell count (% in each B cell cluster) |
|---|---|---|---|---|---|
| 382-65 | B | 6384 (37.6%) | GC B | 496 (10.9%) | 300 (60.5%) |
| | CD4+ T | 9080 (53.5%) | LNPC | 127 (2.8%) | 115 (90.6%) |
| | CD8+ T | 1048 (6.2%) | MBC | 2965 (65.4%) | 149 (5.0%) |
| | NK | 125 (0.7%) | Naïve | 943 (20.8%) | 2 (0.2%) |
| | Monocyte | 251 (1.5%) | | | |
| | pDC | 98 (0.6%) | | | |
| 382-67 | B | 6295 (44.7%) | GC B | 880 (14.8%) | 599 (68.1%) |
| | CD4+ T | 5754 (40.9%) | LNPC | 173 (2.9%) | 143 (82.7%) |
| | CD8+ T | 1567 (11.1%) | MBC | 3889 (65.3%) | 3 (0.1%) |
| | NK | 198 (1.4%) | Naïve | 1011 (17%) | 0 (0%) |
| | Monocyte | 205 (1.5%) | | | |

| | | | | | |
|---|---|---|---|---|---|
| | pDC | 62 (0.4%) | | | |
| 382-69 | B | 21340 (70.8%) | GC B | 372 (5.8%) | 200 (54.6%) |
| | CD4+ T | 6655 (22.1%) | LNPC | 474 (7.4%) | 415 (87.6%) |
| | CD8+ T | 1562 (5.2%) | MBC | 1970 (30.8%) | 5 (0.3%) |
| | NK | 145 (0.5%) | Naïve | 3585 (56%) | 0 (0%) |
| | Monocyte | 337 (1.1%) | | | |
| | pDC | 92 (0.3%) | | | |
| 382-70 | B | 2831 (16.3%) | GC B | 132 (5.3%) | 61 (46.2%) |
| | CD4+ T | 12329 (71.2%) | LNPC | 28 (1.1%) | 21 (75.0%) |
| | CD8+ T | 1827 (10.5%) | MBC | 1620 (64.7%) | 2 (0.1%) |
| | NK | 123 (0.7%) | Naïve | 723 (28.9%) | 0 (0%) |
| | Monocyte | 141 (0.8%) | | | |
| | pDC | 70 (0.4%) | | | |
| 382-71 | B | 4206 (24.4%) | GC B | 138 (3.7%) | 38 (27.5%) |
| | CD4+ T | 11252 (65.2%) | LNPC | 42 (1.1%) | 33 (78.6%) |
| | CD8+ T | 1512 (8.8%) | MBC | 1768 (47.3%) | 0 (0%) |
| | NK | 102 (0.6%) | Naïve | 1792 (47.9%) | 0 (0%) |
| | Monocyte | 103 (0.6%) | | | |
| | pDC | 78 (0.5%) | | | |
| Combined | B | 41056 (41.9%) | GC B | 2018 | 1198 (59.4%) |
| | CD4+ T | 45070 (47.1%) | LNPC | 844 | 727 (86.1%) |
| | CD8+ T | 7516 (7.8%) | MBC | 12212 | 159 (1.3%) |
| | NK | 693 (0.7%) | Naïve | 8054 | 2 (0.02%) |
| | Monocyte | 1037 (1.1%) | | | |
| | pDC | 400 (0.4%) | | | |

**Supplementary data table S5. Cryo-EM data collection and refinement statistics**

| PDB ID 9E21 and EMD- 47426 | |
|---|---|
| **Data Collection** | |
| **Grid type** | UltrAuFoil gold R1.2/1.3 |
| **Microscope/voltage/detector** | Titan Krios/300 kV/Gatan K3 |
| **Magnification** | 105,000 |
| **Recording mode** | counting |
| **Total dose** | 48.95 e-/Å$^2$/s |
| **Pixel size** | 0.825 Å/pixel |
| **Defocus range** | -0.9 to -2.3 μm |
| **No. micrographs used** | 6,614 |
| **Total particles picked** | 630,579 |
| **Model Validation** | |
| **Composition (#)** | |
| **Chains** | 4 |
| **Atoms** | 3285 |
| **Residues** | Protein: 424 Nucleotide: 0 |
| **Ligands** | NAG: 1 |
| **Bonds (RMSD)** | |
| **Length (Å) (# > 4sigma)** | 0.004 (0) |
| **Angles (°) (# > 4sigma)** | 0.749 (0) |
| **MolProbity score** | 1.39 |
| **Clash score** | 3.40 |
| **Ramachandran plot (%)** | |
| **Outliers** | 0.00 |
| **Allowed** | 3.83 |
| **Favored** | 96.17 |
| **Rotamer outliers (%)** | 0.00 |
| **Cbeta outliers (%)** | NA |
| **Peptide plane (%)** | |
| **Cis proline/general** | 0.0/0.0 |
| **Twisted proline/general** | 0.0/0.0 |
| **CaBLAM outliers (%)** | 2.18 |
| **Data** | |
| **Lengths (Å)** | 59.40, 74.25, 108.07 |
| **Angles (°)** | 90.00, 90.00, 90.00 |
| **Supplied Resolution (Å)** | 3.1 |
| **Resolution Estimates (Å)** | Masked |
| **d FSC (half maps; 0.143)** | 3.1 |
| **d 99 (full/half1/half2)** | 2.2/1.7/1.7 |
| **d model** | 2.0 |
| **d FSC model (0/0.143/0.5)** | 1.4/1.8/3.3 |

| | |
|---|---|
| **Map min/max/mean** | -0.00/2.31/0.03 |
| **Model vs. Data** | |
| **CC (mask)** | 0.78 |
| **CC (box)** | 0.63 |
| **CC (peaks)** | 0.49 |
| **CC (volume)** | 0.78 |
| **Mean CC for ligands** | 0.83 |
